# The self as a generative, teleological, and subjective prior: Mutually-modulated temporal agency

**DOI:** 10.1101/519934

**Authors:** Tomohisa Asai, Shu Imaizumi, Hiroshi Imamizu

## Abstract

How can we certify our existence? Recent theoretical studies have suggested that, in the brain, the *effect* inversely infers the possibility of the existence of the self as the *cause*. While this Bayesian view of the sense of agency is widely accepted, empirical evidence in support of this theory is still lacking. The current study examined outcome-modulated agency in terms of time perception in seven experiments, with a total of 90 participants. It was hypothesized that perceptual generation, not termination, should subjectively infer the existence of the self, even though both include the same stimuli and are driven by the same teleological action. Results suggest support for the hypothesis (Experiments 1 and 2). Participants judged stronger self-agency, detected less delay, or felt shorter duration for auditory generation, compared with termination, which was driven by the same volitional key-press. Furthermore, the main experiment, Experiment 3, focused on temporal probability distributions both for action and outcome (e.g., standard deviations or relative entropy), and concluded that the observed contrast in onset/offset agency indicates a mutual or bidirectional relationship between cause and effect only during agentive action, characterized by active (i.e., teleological) generation. Finally, the concurrent theoretical models for volitional action-modulated time perception are discussed on the basis of the suggested *triplet* (generativity, teleology, and subjectivity) dimensions of agency.

**Extended abstract:** The sense of agency refers to the subjective experience of generating one’s own actions (Gallagher, 2000). Within the past decade, research on this subject has increased, as increased understanding of human agency could be a window into knowing how we are aware of ourselves, and possibly allowing for a better understanding of mental illnesses, like schizophrenia. However, “free will,” “self-consciousness,” and/or “volition” are concepts that are always difficult to empirically investigate (Frith & Haggard, 2018). Researchers do not know exactly what occurs in our sensorium during “agentive action.” However, three conceptions could represent genuine dimensions of agency: generativity, teleology, and subjectivity (Haggard, 2019).

Recent theoretical studies have suggested that, in the brain, the *effect* inversely infers the possibility of the existence of the self as the *cause*. This Bayesian view of the sense of agency may provide an explanation for classical statements, such as “ego cogito, ergo sum (I think, therefore I am),” “cogito cogito, ergo cogito sum (I think that I think, therefore I think that I am),” “I move, therefore I am,” or “I predict, therefore I am.” (e.g., Corlett, 2017). All of these theoretical claims implicitly refer to the Bayesian “inverse” probability or inference (e.g., “thought as action”), where the effect (perception) retrospectively postdicts the cause (action), as well as the action predicting the following perception (Figure 1, including generativity, teleology, and subjectivity as triplet dimensions of agency). This bidirectional temporal relationship between action and perception can be empirically examined in terms of the “certification of the self” or *self as prior*.

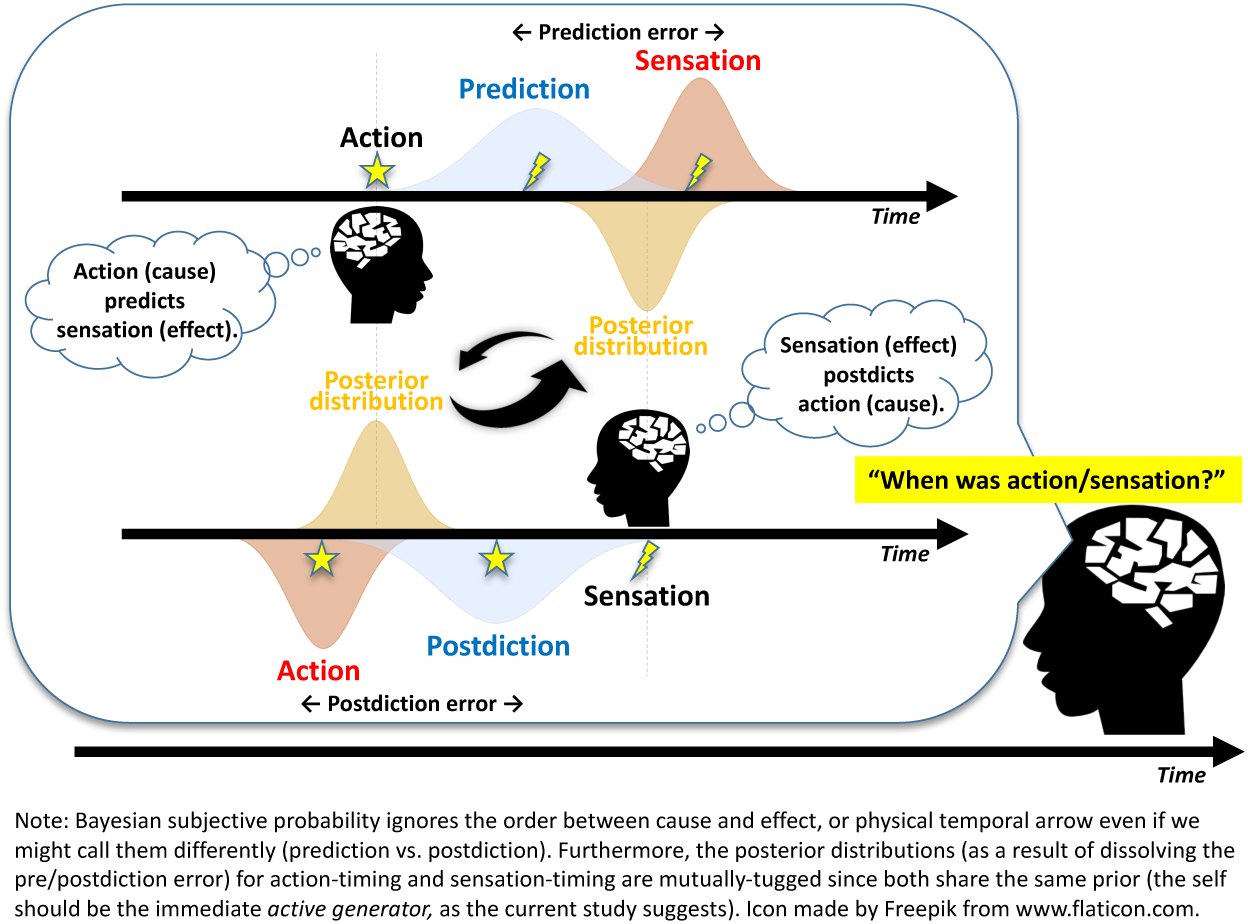

Figure 1.
The theoretical hypothesis of cross-dependency between action and sensation in time.
Note: Bayesian subjective probability ignores the order between cause and effect (i.e., the physical temporal arrow), even if we make a distinction between the two (prediction vs. postdiction). Furthermore, the posterior distributions (as a result of dissolving the pre/postdiction error) for action-timing and sensation-timing are mutually-tugged since both share the same prior (i.e., the self must be the immediate active generator, as the current study suggests). Icon made by Freepik from www.flaticon.com.

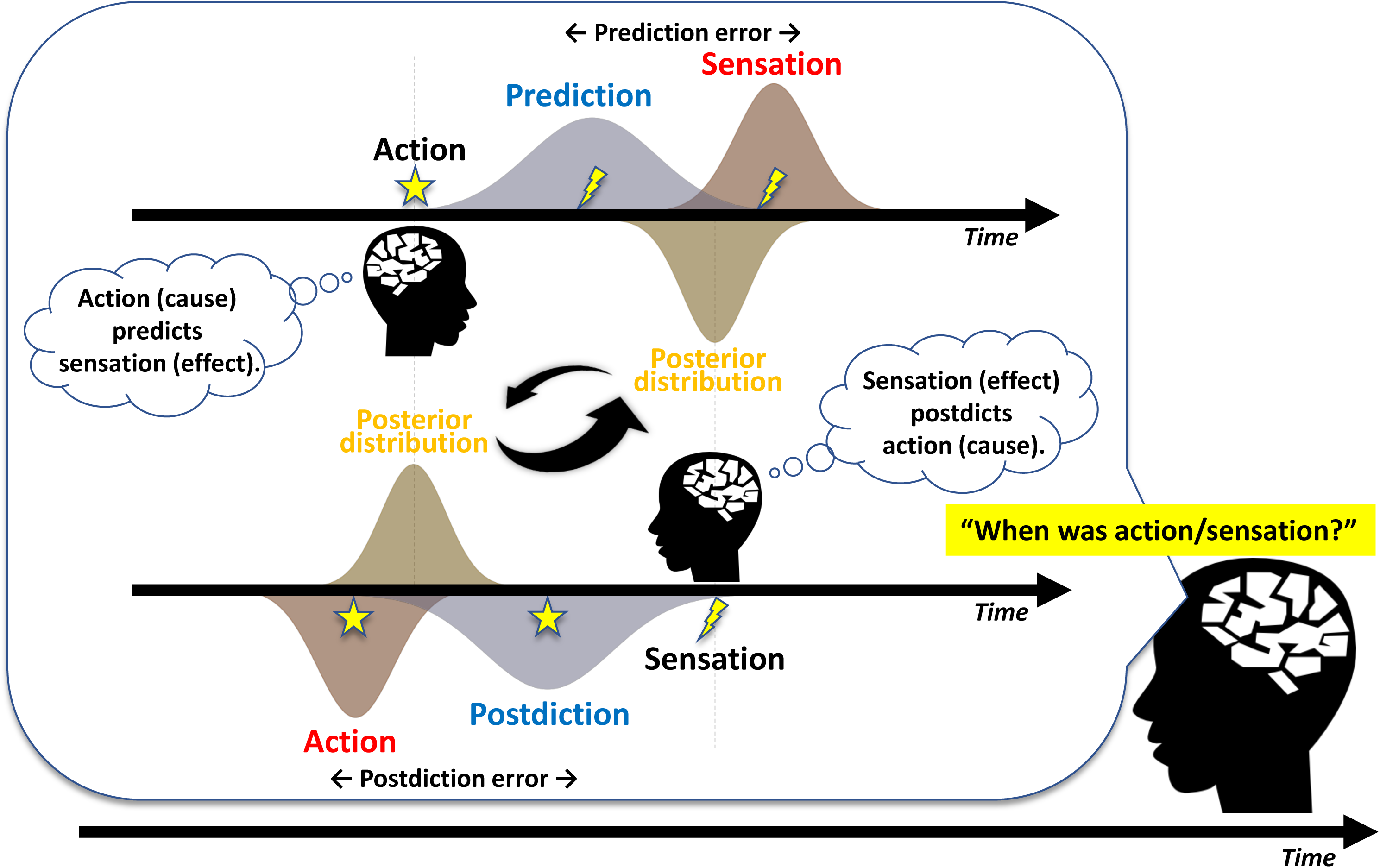

The current study examined effect-modulated agency in terms of time perception in seven experiments, conducted with a total of 90 participants. We hypothesized that perceptual generation, not termination, should inversely infer the self in action, even though both include the same stimuli, and are driven by the same action. Results support this hypothesis (Experiments 1 and 2). Participants judged stronger self-agency, detected less delay, or felt shorter duration for auditory generation, compared to termination, in instances driven by a key-press of his/her own volition, but may have also had different “priors” (i.e., self or other, as a categorical causal structure selection or model weighting). Furthermore, the main experiment, Experiment 3, focused on probability distributions both for action and outcome on the same temporal axis (e.g., standard deviations or relative entropy), and concluded that this contrast in onset/offset agency indicates the mutual or bidirectional relationship between cause and effect only during agentive action, characterized by the active generation (Figure 2). In summary, results indicate that the self is learned as a prior to *being an independent agent* that can immediately cause actions to occur within the environment merely through one’s own volition. This interpretation is highly congruent with the literal meaning of the sense of agency and also with the theory of the triplet dimensions of agency.

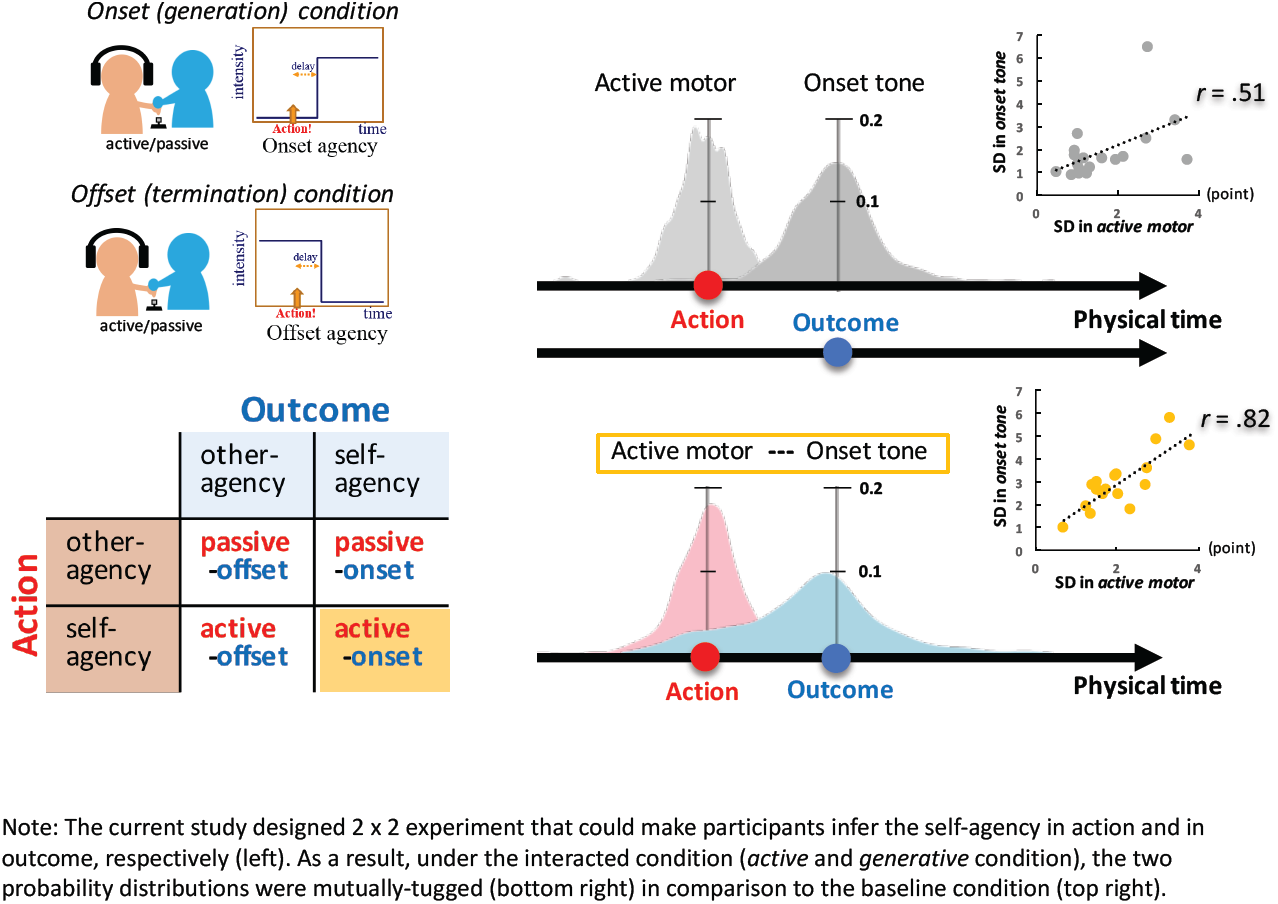

Figure 2.
Summarized results across Experiments 1(A,B,C) and 2(A,B,C).
Note: Asterisks represent the significant differences only between onset and offset conditions. Dashed arches represent the main effect of on/offset condition. Error bars represents ± 1 SE. ** p < .01, * p < .05, † p < .10, see Table S2 for full ANOVA results.

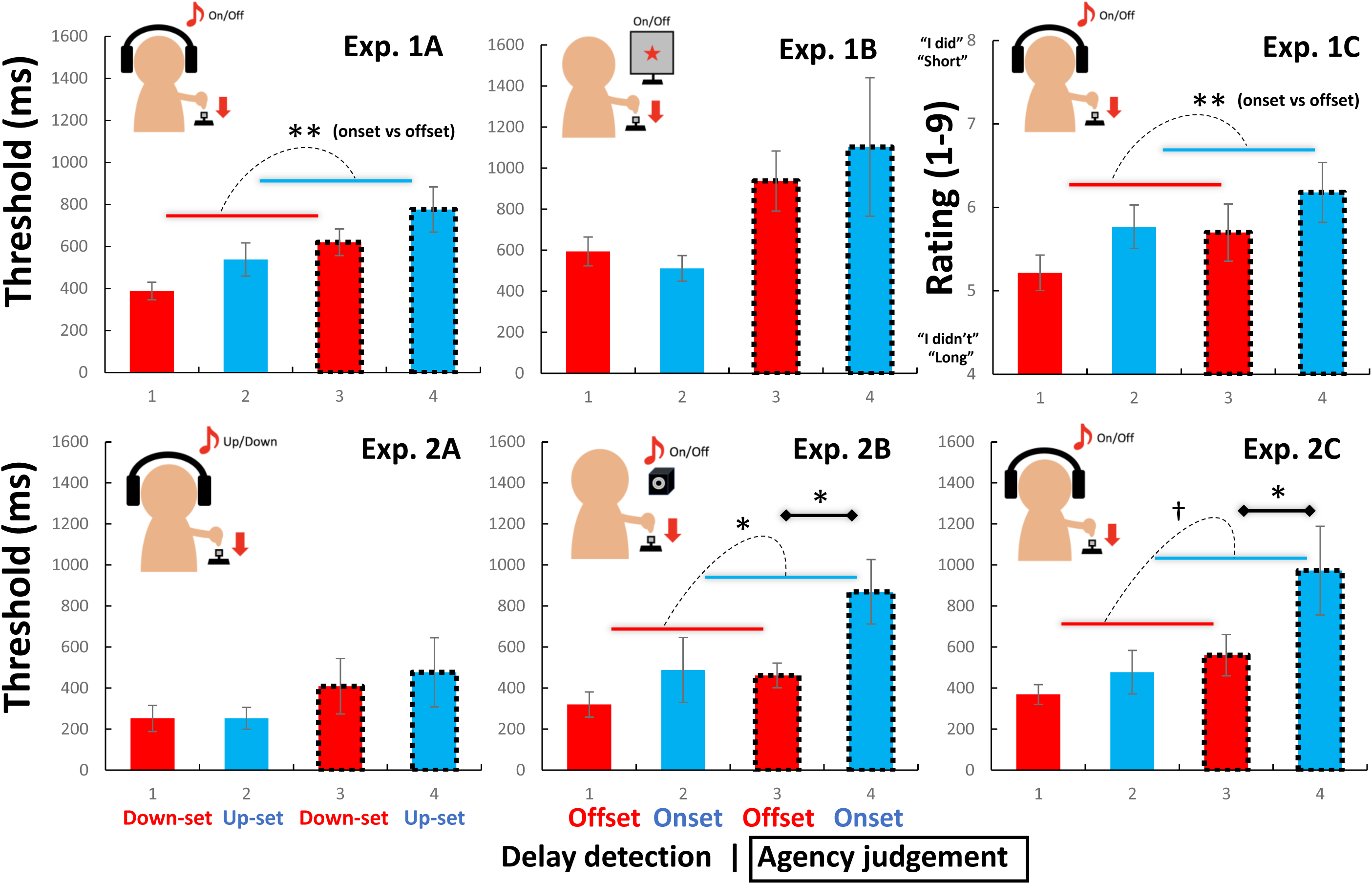

We believe that the current study successfully observed the previously invisible relationship between action and perception in favor of advanced analysis, based on information theory, computational neuroscience (Figure 3), and theoretical discussions.

Figure 3.
The motor, tone, and intentional binding effect in Experiment 3.
Note: Dashed arches represent the main effects of factors. Error bars represents ± 1 SE. ** p < .01, * p < .05, † p < .10, see Table S2 for full ANOVA results.

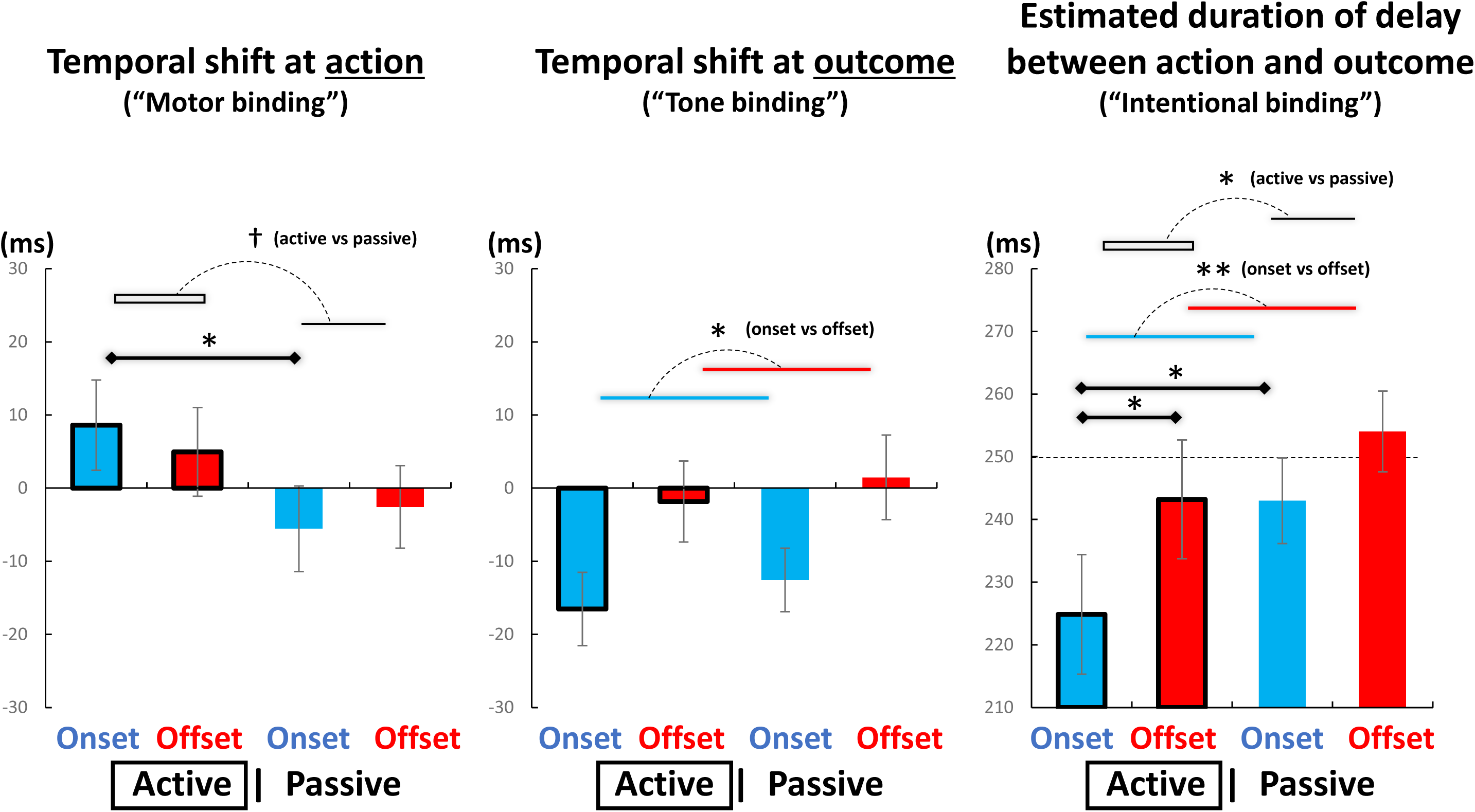

This manuscript has not been published or presented elsewhere in part or in entirety. There are no conflicts of interest to declare.

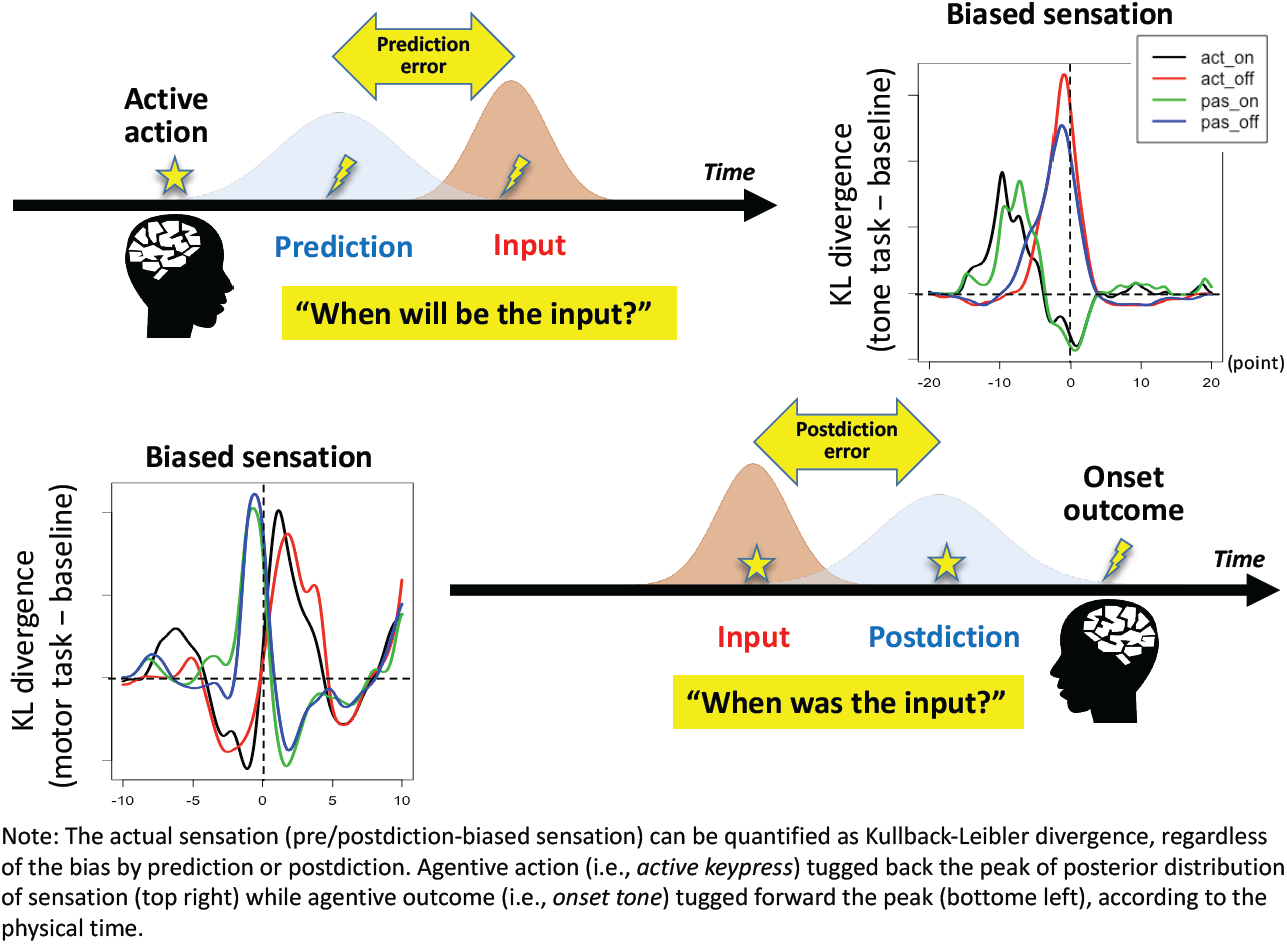

References
Corlett, P. I Predict, Therefore I Am: Perturbed Predictive Coding Under Ketamine and in Schizophrenia. Biological Psychiatry, 81, 465–466 (2017).
Frith, C. D., Haggard, P. Volition and the Brain - Revisiting a Classic Experimental Study. Trends Neurosci. 41, 405–407 (2018).
Gallagher, S. Philosophical conceptions of the self: implications for cognitive science. Trends Cogn. Sci. 4, 14–21 (2000).
Haggard, P. The Neurocognitive Bases of Human Volition. Annu Rev Psychol. (in press)

## Introduction

Recently, agency has become the empirical keyword for researchers who tackle free will, self-consciousness, and/or volition. The shared problem behind these topics can be characterized into three separate dimensions that can be treated as a whole: generativity, teleology, and subjectivity (Haggard, 2019). These dimensions can be further examined by empirical factors. For generativity, the capacity to trigger an event, can be differentiated by the onset (generation) vs. offset (termination) action. For teleology, goal-directedness can be exemplified by the active (goal directed) vs. passive (no goal) action. For subjectivity, the conscious, or probable, inference, can be examined by the interplay between prediction (of outcome) and postdiction (of action), within the temporal arc (see Figure 1 for overview). These issues have been previously examined on an individual basis. However, what makes this “self” problem so complicated (but fascinating) is how difficult it is to understand these separate aspects as a whole, despite the knowledge that the entity must be the agent of action. The current study attempted to show (through behavior-only experiments) that this *triplet* (i.e., generativity, teleology, and subjectivity) is not merely a “working definition,” but rather genuine dimensions that help to both produce and restrain conscious experiences in terms of action and perception (see Figure 4 for an outline of the current experimental protocol).

**Figure 4.**
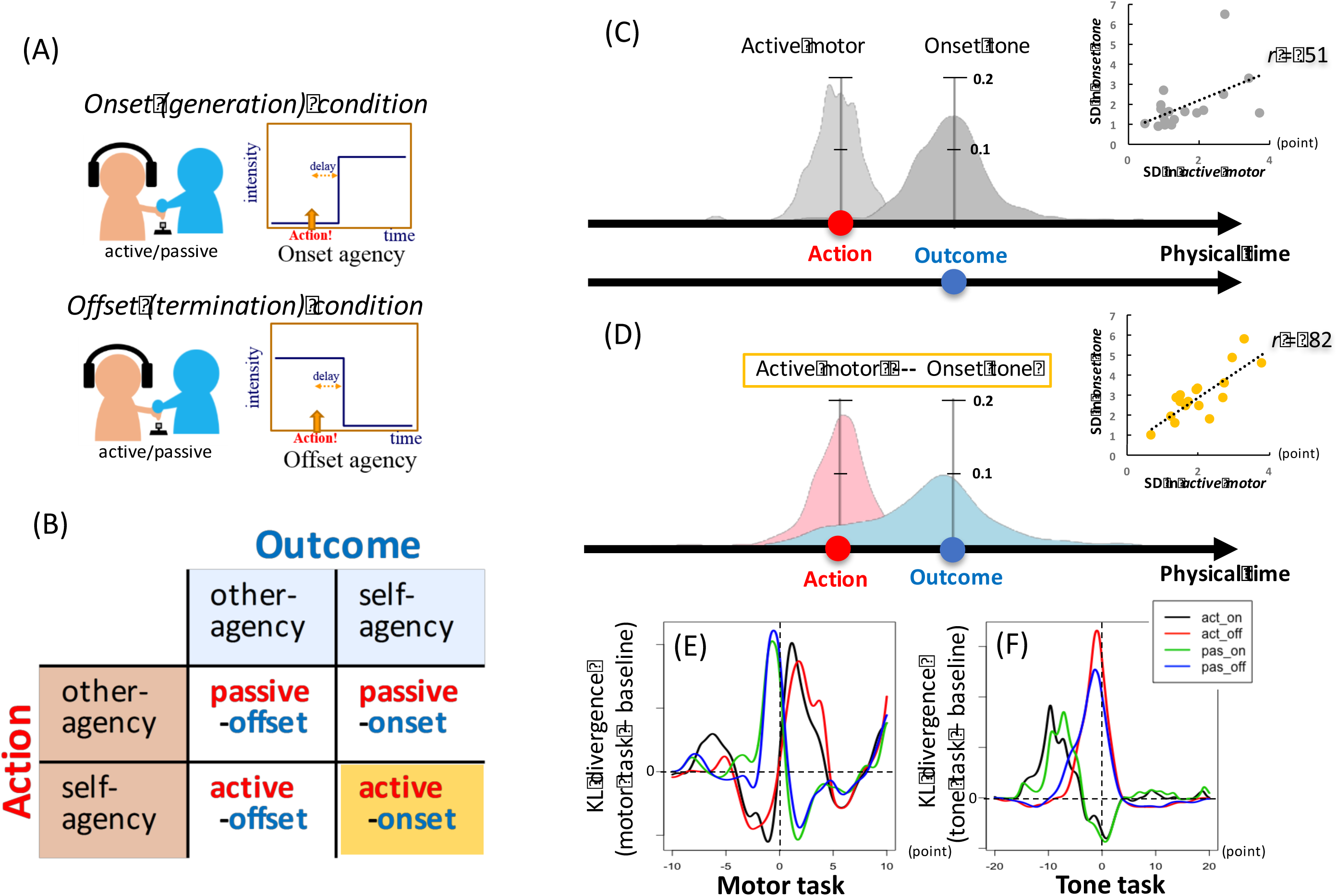
Experimental design (AB), mutually-tugged density plots (CD), and relative peak shifts (EF). Note: Experiment 3 was designed as a 2 x 2 experiment so that participants could infer self-agency in action and in outcome, respectively (AB, see also Figure S1). As a result, under the interacted condition (active and generative condition), the two probability distributions were mutually-tugged (D) in comparison to the baseline (C). Scatter plots suggest the inter-subject correlation (*n* = 18) between SD in active motor and SD in onset tone conditions (see also Figure S8). The peaks were shifted in a Kullback-Leibler divergence from baseline to the experimental task in motor (C) and tone tasks (D), respectively (see also Figure S5).

### Agency as an empirical window

The sense of agency, literally the feeling of *being an agent* in the environment (Crivelli & Balconi, 2017), or more specifically, the subjective awareness of one’s own volitional action (Jeannerod, 2003), has been discussed primarily through the lens of *generating* sensory outcomes. This has often been examined in terms of “the sense that I am the one who is causing an action (Gallagher, 2000)” or “the feeling of making something happen (Haggard, 2017).” However, some papers have presented a more liberal view of agency as “the feeling of control over one’s actions, and their consequences (Voss, Chambon, Wenke, Kuhn, & Haggard, 2017).” This indicates that philosophers and empiricists implicitly believe “agentive action” to mean “generation” or “production.” Yet, no study to date has explicitly examined the essential question of whether the sense of agency only applies to generation, or if termination should also be involved in the perception of *control* (Beck, Di Costa, & Haggard, 2017; Beyer, Sidarus, Fleming, & Haggard, 2018; Borhani, Beck, & Haggard, 2017; Caspar, Desantis, Dienes, Cleeremans, & Haggard, 2016; Kuhn, Brass, & Haggard, 2013; Moore, Lagnado, Deal, & Haggard, 2009). Therefore, the current study compared onset and offset agency in various perceptual, cognitive, and social experimental conditions, in order to elucidate the prerequisites for being an agent in relation to the sensory environment, from a Bayesian perspective (Moutoussis, Fearon, El-Deredy, Dolan, & Friston, 2014).

### Agency in time

When we act, we feel agency. Specifically, the duration between the action and its sensory outcome is perceived as shorter (Haggard, Clark, & Kalogeras, 2002). This temporal binding effect has been considered in various experimental situations as an index of agency (Beyer, Sidarus, Bonicalzi, & Haggard, 2017; Caspar, Christensen, Cleeremans, & Haggard, 2016; Caspar, Desantis, et al., 2016; Cavazzana, Begliomini, & Bisiacchi, 2017; Khalighinejad & Haggard, 2016; Ritterband-Rosenbaum, Nielsen, & Christensen, 2014) c.f.(Dewey & Knoblich, 2014). As a result, we can discuss agency in terms of prediction error (in perception, (Ruess, Thomaschke, & Kiesel, 2017)), contingency (in cognition, (Khalighinejad & Haggard, 2016)), and even responsibility (in social contexts, (Caspar, Christensen, et al., 2016)) within the same temporal domain. Unfortunately, the essential role/meaning is often treated poorly, with an arbitrary or artificial correspondence between action and its outcome presented during the experiment (Buehner, 2012; Desantis & Haggard, 2016). A concrete understanding of the essence of agentive action has yet to be achieved. Previous studies have included many actions, such as single key-pressing (e.g., Sato & Yasuda, 2005), continuous motor control (e.g., Knoblich & Kircher, 2004), or even gesturing (e.g., Daprati et al., 1997). Therefore, the sensory outcomes (e.g., sensory modalities, including vision or audition) depend on the action as the generator (Mifsud & Whitford, 2017; Ruess, Thomaschke, & Kiesel, 2018). Though previous studies have suggested that unnatural (Caspar, Cleeremans, & Haggard, 2015; Ebert & Wegner, 2010) c.f. (Khalighinejad & Haggard, 2016), unreasonable (Takahata et al., 2012), or improbable outcomes (Haggard et al., 2002; Moore & Haggard, 2008) c.f.(Ruess et al., 2017) are less temporally associated (i.e., indicating other-agency), it is necessary to compare between the same sensory stimuli (e.g., tone) and same action (e.g., key-press), but with different meanings or “priors” (Farrer, Valentin, & Hupe, 2013) in terms of self-agency (Moutoussis et al., 2014).

### Action for generation/termination

Roughly half of our actions might work as generators, with the other half acting as terminators. If we do not wait for the spontaneous cessation of generated sensations, we need to terminate that sensation. These sensory onsets and offsets might correspond to neuronal firing in the sensory and motor cortices within our brain (Hubel & Wiesel, 1962). In this sense, generation and termination impulses are routinely coupled throughout our daily lives. This can be conceptualized by turning on/off a radio (auditory sensation) or light (visual sensation) through the same action (i.e., key pressing, (Buehner, 2012) c.f. (Zhao, Chen, Yan, & Fu, 2013)). However, the question still remains as to whether differences exist between the meaning associated with generation and termination. One possible difference may be related to agency, as many previous studies have implied (e.g., agency only in generation, (Khalighinejad, Schurger, Desantis, Zmigrod, & Haggard, 2018; Mifsud & Whitford, 2017)).

Classical ideomotor theory, and other related frameworks, have suggested that action is coupled with perception (for review, see (Gentsch, Weber, Synofzik, Vosgerau, & Schutz-Bosbach, 2016). This means that action is followed by a specific perception of the *effect* of said action. In turn, the perceptual sensation inversely infers the specific action as the *cause*. Here, the question can be rephrased as: What kind of perceptual outcome infers self-oriented action (e.g., volitional key-press)? The literal meaning of the sense of agency (i.e., being an independent agent in the environment) (Crivelli & Balconi, 2017) suggests that agentive action should imply *generation*, as *termination* suggests the presence of others in the world who originally produced the sensory event (e.g., the presence of others, (Beyer et al., 2018; Engbert, Wohlschlager, Thomas, & Haggard, 2007; Moutoussis et al., 2014). Therefore, the contrast between the perceived outcome of generation and of termination is crucial for inversely inferred agency in action, where action is coupled with sensory generation. Surprisingly, no study has examined action as a generator compared to action as a terminator in terms of agency, though the sensory onset/offset mechanism is fundamentally driven by the identical goal-directed action (i.e., teleology).

### Bayesian inference from perception to action

Recent theoretical frameworks of perception, cognition, and action are commonly based on Bayesian “inverse” probability (Baker, Saxe, & Tenenbaum, 2009; Buehner, 2012; Gentsch & Synofzik, 2014; Kording & Wolpert, 2006). In this view, our brain works as a prediction machine, determining the future based upon the past and present. Another conceptualization of this view is the brain as an inference machine, inferring the cause (in the past) from the effect (in the present), regardless of the domains (Lochmann & Deneve, 2011; Miall & Wolpert, 1996; Sharps & Martin, 2002; Uithol & Paulus, 2014). The shared mechanism among these conceptualizations is prediction error. As computational motor control theory suggests (Wolpert, 1997), we learn the internal model for our own actions, and even adapt this model to the external environment. This model can serve as a prior cue to predict, or provide inference of the hidden state (the future or the past), while simultaneously minimizing detected prediction error between the prior and the sensory data available (through either changing the prediction or changing the sensory samples). Accordingly, we can use unexpected (i.e., unpredictable) outcomes to learn about the world and ourselves, by continuously updating our beliefs of “how our sensations are caused” (Brooks, Carriot, & Cullen, 2015; K. Friston, 2010; Stahl & Feigenson, 2015). In this conceptualization, we can see Bayesian inference (i.e., subjectivity), not only from action to perception (i.e., teleology), but from perception to action (i.e., generativity).

The essential question of the current series of experiments (i.e., by what outcome the volitional key-press is inversely inferred) is Bayesian friendly. In predictive coding (K. J. Friston, Stephan, Montague, & Dolan, 2014; Rao & Ballard, 1999), the reliability of sensory information, as well as the sensory input itself, can be predictable. This metacognitive capacity is referred to as the “precision” or “confidence” assigned to various prediction errors that ascend cortical hierarchies (Adams, Stephan, Brown, Frith, & Friston, 2013; Picard & Friston, 2014), to infer the *relative precision* placed on sensory evidence and prior beliefs (Wolpe et al., 2016). Precision is psychologically associated with attentional gain and physiologically with the excitability of neuronal populations reporting prediction errors, resulting in a modulation of perceptual variance. Recently, this view has been proposed as the basis of many neuropsychiatric disorders (Adams, Stephan, et al., 2013; Lawson, Rees, & Friston, 2014; Picard & Friston, 2014; Teufel et al., 2015), in addition to agency-related perceptual or cognitive phenomena (e.g., intentional binding, sensory attenuation, or self-other discrimination (Brown, Adams, Parees, Edwards, & Friston, 2013)), especially in terms of “biased” perceptual variance or precision by priors (Wolpe, Haggard, Siebner, & Rowe, 2013).

### Onset/offset agency

The key purpose of the current study was to temporally examine onset and offset agency. The former can be defined as the subjective feeling of generating an action itself, as well as through a difference in time perception (delay detection, duration estimation, or timing report) (Engbert et al., 2007; Haggard et al., 2002; Makwana & Srinivasan, 2017) c.f. (Farrer et al., 2013) in the sensory event generated by the action. The latter is defined as the feeling of agency for terminating an action and the time perception of the terminated event by the individual’s action. We hypothesized that onset agency is more evident than offset agency if self-agency is rooted in generation (Gallagher, 2000; Haggard, 2017; Jeannerod, 2003; Khalighinejad et al., 2018; Legrand, 2007; Mifsud & Whitford, 2017). Conversely, termination implies the action of others prior to termination, suggesting that the agentive action should entail a special state of consciousness or sensorium to temporally predict agentive outcomes (Bays, Wolpert, & Flanagan, 2005; Picard & Friston, 2014; Walsh & Haggard, 2013). To examine these concepts, seven experiments were conducted in a total 90 participants. Following descriptions of these experiments, the concurrent theoretical models are discussed within the framework of the current results.

## Experiment 1 (A, B, C)

The purpose of Experiment 1 was to simply examine the contrast between onset and offset agency in relation to detection delay. Participants were always required to make a volitional key-press, but this action would cause one of two types of outcomes with a delay, specifically, sensory generation or termination. Following this, participants were required to report subjective feelings of agency and any delay that was detected.

### Methods

#### Participants

A total of 47 healthy individuals (males = 19, mean age = 33.8, standard deviation [SD] = 9.3) participated in Experiment 1 (17, 18, and 12 participants for Experiments 1A, 1B, and 1C, respectively). Participants were recruited from the local community and were paid for their participation. All participants were right-handed and reported normal or corrected-to-normal vision and hearing. Further, all provided written informed consent prior to experiments being conducted. The experiments were conducted in accordance with the Declaration of Helsinki. The protocol of the present study was approved by the local ethics committee.

#### Apparatus

A standard LED monitor, headphones, and a keyboard were used for the instruction, stimulus presentation, and key-pressing. The visual or auditory stimuli were controlled by Hot Soup Processor 3.3 (Onion Software) installed on a Windows computer. A simplified illustration of the apparatus is shown in Figures 1 and 2.

#### Procedure

The basic procedure of the experiment followed those outlined by previous studies (see for review, (Haggard, 2017; Moore & Fletcher, 2012)), with participants required to press a key with their right index finger as the action. Participants were instructed that they could press the key at their own pace after a cue in each trial. Each experiment had 2 x 2 task design: agency judgement/delay detection blocks x onset/offset conditions. In the agency judgement block, participants in Experiments 1A and 1B were instructed to report self- or other-agency by a Two Alternative Forced Choice (2AFC) response. In other words, participants had to report the intuitive feeling about the action-outcome, specifically whether they felt they were responsible for the sensory event (i.e., as the generator or terminator, see below for onset/offset condition). Conversely, in the delay detection block, participants were instructed to report if a delay was inserted between their key press and the outcome. In Experiment 1C, rather than a 2AFC, participants rated level of agency or delay on a 9-point Likert scale, with a score of 9 representing *I caused the outcome* or *The delay was very long*, and 1 representing *I did not cause the outcome* or *No delay*)*short)* (Sato & Yasuda, 2005). The order of the two blocks (i.e., agency judgement and delay detection) were counterbalanced among participants. They were explicitly instructed that they did not need to make an agency judgment on the basis of delay, so that they could intuitively judge in the agency block. This was also true for the delay block, where they could report their delay detection, regardless of agency (Asai & Tanno, 2007, 2008, 2013).

Each block included two further conditions (Figure 1). Under the onset condition, a trial started with silence. When participants made a key press, an auditory tone (Experiments 1A and 1C) or a visual symbol (Experiment 1B) were presented after a short delay. The stimuli continued to be presented after participants’ response for agency or delay detection by pressing a corresponding key. Once response was made, the stimulus gradually disappeared (i.e., faded out), unlike when the stimulus was generated. Participants used three keys: one for the action, and the other two keys for the response. Under the offset condition, the stimulus gradually appeared (i.e., faded in) before a cue for action, and continued to be presented. Once participants pressed the key as the action, the stimuli disappeared instantly or with delay (depending on condition). After this, participants responded about agency or delay. The two conditions were presented randomly. Furthermore, the tones/symbols were randomly varied in each trial (tone ranged from 500 to 1000 Hz, with 100 Hz step; symbols were a solid circle, square, triangle, star, rhombus, and double circle), so that participants could not bind block or condition with a specific stimulus. This also served to prevent carry-over effects across trials (e.g., the star is for self-agency). The amplitude of the auditory tone was approximately 70 dB SPL. The visual angle of the symbol was approximately 5 degrees. Both the auditory and visual stimuli were easily discernable for all participants.

The delay inserted between the action and the outcome was one of five delay conditions (0, 100, 300, 700, and 1500 ms). These conditions were randomly presented in each agency/delay block and onset/offset condition. Each delay condition was repeated 10 times. Therefore, participants had 100 trials (5 delay conditions x onset/offset conditions x 10 repetitions) in both the agency delay blocks. Prior to commencement of the experiment, participants were briefly trained to help them get accustomed to the device and procedures.

#### Analysis

The data obtained in the current study were analyzed by R 3.4.2 (R Core Team, 2017). We also used the *ggplot2* package (Wickham, 2009) to visualize results. First, each participant’s binary raw responses were fitted with a cumulative normal distribution as a psychometric function, using a maximum-likelihood procedure. This probit analysis revealed 50%-thresholds in each block/condition for each participant, as shown in Figure S2. The mean raw reports were also summarized, as well as the mean conditional thresholds (or delay condition-collapsed average for Experiment 1C) as the main results. The full analysis of variance (ANOVA) tables for this main result are also provided as a supplementary table (Table S1).

### Results & Discussion (Experiment 1)

Figure 2 (top) is the summarized results across three experiments (Experiments 1A, 1B, and 1C). The threshold (1A and 1B) for a feeling of agency or detection of delay was determined by the fitted psychometric curve for each participant (see Analysis, Figure S1 and Figure S2). When examining feelings of agency, a longer delay above threshold suggests other-agency and vice-versa. When examining detection of delay, a longer delay above threshold suggests that the delay was detectable. Measures of continuous ratings (1C) were similar for both conditions, but more precise, as these measures provided an estimation of magnitude. Participants rated subjective self- or other-agency, or the subjective duration between the action and the outcome (see Procedure). The mean ratings (collapsing delay-conditions) for each participant were compared among four conditions to keep a statistical congruence with Experiments 1A and 1B (see Figure S1 for detailed results).

#### Positive bias of agency in relation to delay

As suggested in previous studies, we often observed a positive or self-serving bias in agency judgement (Asai & Tanno, 2007, 2013; Beyer et al., 2017; Miyazaki & Hiraki, 2006; Takahata et al., 2012). A significant main effect of agency/delay block in Experiments 1A and 1B (but not in 1C for rating, as there was no absolute temporal reference) suggests a positive bias [*F*(1,16) = 9.325, *p* = 0.0076 for Experiment 1A, *F*(1,17) = 5.441, *p* = 0.0322 for Experiment 1B, see Table S1 for the full ANOVA table). This means that, even when we can detect a small bias or incongruence between the action and the outcome (e.g., delay), we tend to allow that small “prediction error” to be attributed to ourselves to update our internal model about the self and the environment (Adams, Stephan, et al., 2013). The sensory attenuation suggests that the volitional action (e.g., keypress) should produce the sensory outcome (e.g., tone) without delay (Bays et al., 2005; Blakemore, Frith, & Wolpert, 1999). This learned prior recalibrates the perceived temporal order between the action and outcome (Stetson, Cui, Montague, & Eagleman, 2006), where the exposed delay as prediction error can change our attentional gain to maintain our belief of no-delay (i.e., active inference in predictive coding). In this sense, the other option of updating belief (my action might produce the sensory outcome with substantial delay) seems to be more difficult than reversing the cause and effect in our perception, especially when priors are strong in precision (e.g., faces (Shipp, Adams, & Friston, 2013)).

#### Onset and offset agency

Regarding the difference between onset and offset conditions as the primary outcome measure of interest in the current study, the main effect of on/off condition was significant in Experiment 1A (*F*(1,16) = 12.013, *p* = 0.0032), an effect that was also seen in Experiment 1C, but not in Experiment 1B. This indicates that, in the onset condition, where participants were made the generator of the sensory event, the temporal threshold for delay detection, and also for agency judgment, was elevated, when compared with the offset condition as the terminator of the stimulus. This is because participants perceived a shorter duration and more agency in the onset condition than in offset condition (Experiment 1C, *F*(1,11) = 30.989, *p* = 0.0002). This onset/offset contrast was not due to the aftereffect of the offset condition, where participants might perceive illusional stimuli, even after offset timing, since this does not explain the reduced threshold in Experiment 1A, and because the visual onset/offset did not modulate both thresholds in Experiment 1B (also see Experiment 2C for delay-parameter independence). Rather, this result indicates that action-driven onset is temporally shifted toward the timing of the action or our conscious resolution of time may be altered, but only in onset situations (i.e., for waiting for a self-generated outcome in “compressed time” or in a “feedback mode,” see General discussion for models).

#### The outcome in modalities

In contrast to Experiments 1A and 1C, which used auditory outcomes, the difference between onset and offset conditions (i.e., strong positive bias for self-agency and enhanced delay detection as a generative agent) was not found in Experiment 1B, which used visual outcomes [*F*(1, 17) = 0.115, *p* = 0.7385]. This visual inferiority is consistent with previous findings on the other perceptual phenomena related to agency. For instance, the intentional binding effect, whereby subjective timings of voluntary action and its sensory outcome attract each other ((Haggard et al., 2002), also see Experiment 3), is weakened for a visual outcome relative to an auditory outcome (Ruess et al., 2018). Sensory attenuation, another agency-related perceptual modulation, whereby individuals are likely to perceive decreased intensity of sensory events caused by themselves, than by others or the environment, is robust in tactile (Blakemore et al., 1999) and auditory (Weiss, Herwig, & Schutz-Bosbach, 2011) domains, while “visual” attenuation is unlikely to occur (Schwarz, Pfister, Kluge, Weller, & Kunde, 2018).

To date, the mechanisms underlying modality specificity in intentional binding and sensory attenuation remain unclear. A potential explanation is that our motor system is more behaviorally (Repp & Su, 2013) and neurally (Hove, Fairhurst, Kotz, & Keller, 2013; Jancke, Loose, Lutz, Specht, & Shah, 2000) coupled to the auditory system than to the visual system. For instance, manual synchronization to rhythmic auditory stimuli has been known to show better performance than manual visual rhythmic synchronization (Repp & Su, 2013). Extending these notions within an ideomotor framework (e.g., (Gentsch et al., 2016)), our bodily actions may be more strongly related to auditory events in the external environment, where sounds are typically generated by manual action, gait, or vocalizations. Auditory events serve an essential function in survival (e.g., warning and communication). Considering this, sense of agency can be modified by generation and termination of events in the environment, especially when our agentive action is within the auditory domain.

## Experiment 2 (A, B, C)

Though the three experiments in Experiment 1 suggested a contrast between onset and offset agency (i.e., the self as the generator), follow-up experiments were necessary. First, in Experiment 2A, a comparison was made between up-/down-set, rather than onset/offset, in an effort to confirm the “generation” effect. Additionally, tone localization was examined in Experiment 2B to explore an outcome “in the environment” (Engbert et al., 2007; Hon, Seow, & Pereira, 2018), not within ourselves. Finally, delay-parameter independence was confirmed in Experiment 2C in an effort to increase the generalizability of the findings.

### Methods

A total of 23 healthy individuals (males = 5, mean age = 31.2, SD = 8.9) participated in Experiment 2 (8, 8, and 7 participants for Experiments 2A, 2B, and 2C, respectively). Procedures were almost identical to those presented in Experiment 1A (i.e., delayed auditory event for agency judgement or detection of delay), except for the following (also see Figure 2 for comparison). In Experiment 2A, a shift in tone volume (either up or down) was the outcome, rather than on/off. The initial volume was 60 dB SPL, but this was raised via keypress, with a delay, to 70 dB SPL in Up condition, and lowered to 50 dB SPL in the Down condition. The apparatus and other procedures were identical to Experiment 1A. In Experiment 2B, the tone was presented through a standard speaker, instead of headphones. The speaker was located approximately 30 cm in front of participants, but visually occluded from the point where they were instructed to look at the monitor for the action. In Experiment 2C, the range and steps of delay were altered, so that the longest delay was 1200 ms (0, 200, 500, 800 and 1200 ms), rather than 1500 ms (0, 100, 300, 700 and 1500 ms) that was utilized in Experiment 1A, in order to avoid a dependency on the specific experimental parameter (i.e., the range of delay in this case) that is used to fit the psychometric function. The task itself was also identical, where participants were required to press a key with their right index finger as the action. Participants were instructed that they could press the key at their own timing after a cue in each trial, under a 2 x 2 task design: agency judgement/delay detection blocks x onset(up)/offset(down) conditions.

### Results & Discussion (Experiment 2)

The summarized results across the three experiments (Experiment 2A, 2B, and 2C) are shown in Figure 2 (bottom). Threshold was determined in the same manner, and other procedures were almost identical to those described in Experiment 1A, except for the comparison between up-/down-set in Experiment 2A, the tone through the speaker in 2B, and the changed delay-parameter in 2C.

#### Not up-set regulation, but generation is necessary for agency

The onset/offset manipulation in Experiment 1 included up-/down-set regulation. It was unclear if generation (i.e., to produce something from nothing in terms of sensation) is actually necessary for temporal binding and agency. However, the results of Experiment 2A do not provide support for this possibility, as there was no significant up-/down-set condition effect or agency/delay block effect observed [*F*(1,7) = 2.805, *p* = 0.1379 for agency/delay effect, *F*(1,7) = 0.821, *p* = 0.3950 for up-/down-set effect).

#### Generation outcomes in the environment

The results of Experiment 2B essentially replicated the results of Experiment 1A, where a significant main effect of onset/offset was revealed. In addition to this, the interaction was also significant [*F*(1,7) = 6.365, *p* = 0.0396) and a simple main effect of onset/offset condition in agency block was also identified. Though a fair comparison between Experiment 1A and 2B is difficult (e.g., due to a difference in sample size), agency might be elicited more strongly by onset outcome in the external environment than from immediate proximity to one’s own body, suggesting a need for self to generate outcomes in the world (Engbert et al., 2007) c.f. (Hon et al., 2018).

#### Parameter independence

A merit of the comparison of thresholds is that, by definition, they are parameter-free. As can be seen by the raw results in Figure S1, the significant difference may only be observed under specific delay conditions, where the delay parameter was decided a priori by experimenters. Therefore, such statistical significance in a specific condition has no essential meaning. Indeed, the estimated thresholds in Experiment 2C was almost identical to Experiment 1A (delay parameters were 0, 200, 500, 800, and 1200 ms for Experiment 2C, and 0, 100, 300, 700, and 1500 ms for Experiment 1A), with statistical results basically following suit (although the interaction was significant in Experiment 2C; *F*(1,6) = 8.723, *p* = 0.0255).

In summary, results of Experiments 1 and 2 generally suggest that time perception and feelings of agency about the outcome are modulated when the action is served as a generator; although results are still not conclusive about the outcome modality (1B), report methodology (1C), up/down regulation (2A), tone localization (2B), or delay parameters (2C). Especially agency-specific (non-delay) modulation might be observable in certain situations, as suggested by Experiments 2B and 2C. However, the primary concern should be to determine whether time is biased for action or for outcome. All experiments to this point utilized relative comparison between onset and offset conditions, as there is no definitive answer to thresholds or ratings. The well-known Libet’s task (i.e., clock reading paradigm)(Haggard et al., 2002) is, in this sense, tricky, but useful in making an absolute comparison (as error) according to physical time flow. Especially, to examine bidirectional entrainment or “mutually-biased” hypothesis.

## Experiment 3

The purpose of Experiment 3 was to reexamine the onset-specific modulation of time perception in a clock reading paradigm. An advantage of this task is that it can allow us to examine the absolute report for each timing (action or outcome) from participants. This is especially useful, since we can also examine sensory reliability from the variance (Wolpe et al., 2013) in its distribution plot. Though the experiments performed so far have required participants to perform the same action (e.g., volitional keypress), in this final Experiment 3, the factorial interaction between “agentiveness” in action (active/passive motor) and that in outcome (onset/offset tone) was a special target of interest (i.e., the interaction between teleology and generativity). The mutual effect between these two concepts was hypothesized, where sensory precision was also examined in terms of Bayesian integration (i.e., subjectivity).

### Methods

#### Participants

A total of 20 healthy volunteers (males = 7, mean age = 29.0, SD = 8.5) participated in Experiment 3.

#### Procedure

The onset/offset procedure was essentially the same as presented in Experiments 1 and 2, where participants were required to press a key so that the tone was either generated or terminated with a delay (in this case, always 250 ms). However, participants were also instructed to attend to a clock on the display, ranging from 5 to 60, in intervals of 5 (Figure S1). A single hand rotated, clockwise, with a period of 2560 ms (starting at a random position in each trial). Participants reported the time of the action or the outcome (see below for detailed conditions) using a keyboard at the end of each trial (Haggard et al., 2002).

A total of 12 conditions were administered for each participant. The baseline-tone block included two conditions: onset tone and offset tone conditions were randomly presented (10 repetitions for each). Participants reported the time the tone appeared in the onset condition, or when the tone disappeared in the offset condition, without any prior action (the onset/offset was random between 1500 and 2500 ms after each trial started). The baseline-motor block included 2 conditions: active and passive motor conditions were further blocked in a random order (10 repetitions for each). Participants reported the time of their own self-paced key press in the active condition or of the experimenter-forced key press in the passive condition, where the experimenter pressed the participants’ index fingertip to depress the key. The active or passive motor action in this block did not result in any sensory outcome. One of the three experimenters was randomly assigned to each participant to avoid an effect of a specific experimenter (e.g., preference for specific timing of key-press).

These control conditions were compared to the experimental conditions below. The active action-outcome block included 2 x 2 conditions, presented in a random order (25 repetitions for each): onset/offset and reporting target (action or outcome). After the self-paced key press, a tone was generated in the onset condition or was terminated in the offset condition, with a 250 ms delay. Since participants were not aware of the reporting target until the answering phase at the end of the trial, they had to memorize two timings but reported just one of the two (or retrieved a specific timing when required). Similarly, the passive action-outcome block included 2 x 2 conditions presented in a random order (25 repetitions for each): onset/offset and reporting target (action or outcome). In this block, however, the key press was forced by the experimenter, in a manner similar to the passive baseline-motor condition (see Figure S4 for design).

On each trial, the clock stopped randomly 1500–2500 ms after the outcome. Participants first completed two control blocks in a random order. After this, they also completed two experimental blocks in a random order. Other procedures and experimental settings were identical to those presented in the description of Experiment 1A. The duration of Experiment 3 was approximately 90 minutes, including a rest period.

#### Analysis

Participants’ raw estimation errors (i.e., the difference between the actual and reported time of the event) are shown in Figure S3 and Table S2, where trials with outliers greater than ± 10 points (= 426 ms) of the mean were removed from each participant’s dataset. Two participants were excluded from analysis as the mean of the baseline condition (baseline-motor or baseline-tone) was greater than ± 2 SD of the group mean. For the binding effect, mean estimation errors in the baseline condition (motor or tone) were subtracted from the corresponding action-outcome conditions (active/passive x onset/offset, respectively) to obtain motor binding, tone binding, and their composite intentional binding measures (see Figure S4). Full ANOVA tables for these main results are provided in Table S1.

Additionally, to test the mutual effect between action and outcome, distributions, as well as SD (and mean) were analyzed by Kullback-Leibler divergence (KLd), inter-subject correlation, hierarchical clustering (e.g., (Ais, Zylberberg, Barttfeld, & Sigman, 2016)), and AI-based Bayesian network modelling (McNally, Mair, Mugno, & Riemann, 2017) (Glymour, 2003). For this, *LaplacesDemon* package (Statisticat LLC, 2016) in R was added to calculate KLd, *TSclust* (Montero & Vilar, 2014) to make a cluster dendrogram, and *bnlearn* (Scutari, 2010) and *Rgraphviz* (Hansen et al., 2017) to compute Bayesian networks and to visualize them as a directed acyclic graph using a tabu search algorithm.

### Results & Discussion (Experiment 3)

The summarized results are shown in Figure 3 (descriptive statistics are provided in Figures S3, S4, S5, S7, and Table S2). Motor binding, tone binding, and the intentional binding effects were individually plotted and discussed, as previous studies have suggested different mechanisms between motor and tone binding (e.g., Wolpe et al., 2013). We also examined the distribution of participants’ raw report of timing (error) as sensory reliability (Figure 4) in order to identify what was modulated in the primary experimental condition.

#### Motor binding

The left side of Figure 3 depicts the motor binding effect, the subtraction between the mean estimation errors for motor timing in baseline-motor conditions (active/passive), and the corresponding action-outcome conditions (active/passive x onset/offset). Previous studies have argued that motor binding can be described both as a postdictive, inferential process, and as a predictive process (Moore & Haggard, 2008). The current results of motor binding, in which the interaction was significant [*F*(1,17) = 0.0166, *p* = 0.0207] suggests that motor timing in the active-onset condition was perceptually shifted toward the outcome tone, in comparison with that in passive-onset condition. This active motor effect, in which the main effect was also marginally significant, supports the notion of predictive modulation (e.g., a pre-activation account (Waszak, Cardoso-Leite, & Hughes, 2012)), where the active motor action is temporally-predictive of the outcome, compared with the passive motor action. Participants could prepare for the outcome, even when they were urged to press the key by themselves. Conversely, this modulation was also postdictive (Moore et al., 2009) in terms of the onset-specific effect. Though the onset or offset of tones temporally followed the motor action, this postdictive difference retrospectively, or inversely, modulated the perceived timing of the motor action (though participants were able to determine the condition, onset/offset, as this was presented during the motor action). The comparison between motor and tone binding is also interesting for contrasting results as follows.

#### Tone binding

The middle section of Figure 3 depicts the tone binding effect, the subtraction between the mean estimation errors for tone timing in baseline-tone conditions (onset/offset), and the corresponding action-outcome conditions (active/passive x onset/offset). Compared to motor binding, the result of tone binding was simple, with a significant main effect of onset/offset [*F*(1,17) = 11.379, *p* = 0.0036]. The onset timing of tone was perceptually shifted back to the action, regardless of its active/passiveness. This modulation should be postdictive, since the outcome (onset/offset) solely had an impact (Desantis, Hughes, & Waszak, 2012) c.f. (Waszak et al., 2012). The result of this tone binding might be a good replication of Experiments 1 and 2, but has advantages in theoretical development. The previous experiments suggested that the threshold in delay detection (or in agency judgment) was elevated in the onset condition (Experiments 1A, 2B, and 2C), meaning the less detectable interval between the motor action and the onset of tone. The direct impression (e.g., magnitude estimation) also suggested a shorter duration between motor action and onset condition (Experiment 1C). Though these results imply a binding effect, they were explicitly not tested (i.e., motor action was always active in Experiments 1 and 2). The current result helped to elucidate this effect. The onset-specific modulation of outcome timing is not depending on whether the motor action is active or passive. As long as the motor action, and its outcome (onset tone), is coupled (Engbert, Wohlschlager, & Haggard, 2008; Khalighinejad & Haggard, 2016), or in other words, as long as the tone is generated by the motor action, the perceived onset timing would shift back. We can also see the result of so-called intentional binding as follows.

#### Intentional binding

The right section of Figure 3 depicts the intentional binding effect, the further subtraction between the mean estimation between in action timing (from motor binding), and in outcome timing (from tone binding) under each condition (active/passive x onset/offset) in reference to the actual interval between them (i.e., 250 ms). As expected, from motor and tone binding, the estimated duration between action and outcome was significantly shorter only under the active-onset condition, demonstrating a significant interaction [*F*(1,17) = 6.041, *p* = 0.0250]. Since intentional binding is the composite score of motor (active/passive marginally significant main effect) and tone (onset/offset significant main effect) binding (left and middle panels of Figure 3), the interaction presented here (right panel of Figure 3) could be significant as a result, in addition to the main effects of active/passive condition [*F*(1,17) = 4.635, *p* = 0.0460] and of the onset/offset condition [*F*(1,17) = 9.925, *p* = 0.0058], while the specific effect (i.e., the simple main effect) was observed only for the active-onset condition. This interaction between action and outcome is worth of further examination. Though the action temporally preceded the outcome, action timing (active condition) was also inversely affected by the outcome (onset condition). The mean, distribution, and variance of this potential mutual coupling was further examined.

#### Sensory uncertainty in time perception during agentive action

A previous study examined increased variance in reports of outcome (SD) as an index of sensory unreliability or uncertainty (Wolpe et al., 2013), as well as the shift of the mean, in terms of Bayesian cue integration theory. According to this theory, the perceived action timing should become uncertain due to the given outcome (and vice versa), since two events (in this case, active motor action and onset outcome) are associated. The no-delay assumption as a prior in self-generated sensory outcomes can weight distributions on the basis of each reliability (see also Discussion section of Experiment 1 and the theorized schematics in Figure 5). As previous experiments have suggested, due to the elevated threshold observed during generative action (e.g., Experiment 1A), time perception appears unreliable in onset agency because the uncertainty between the action and the outcome might be linked to one another.

**Figure 5.**
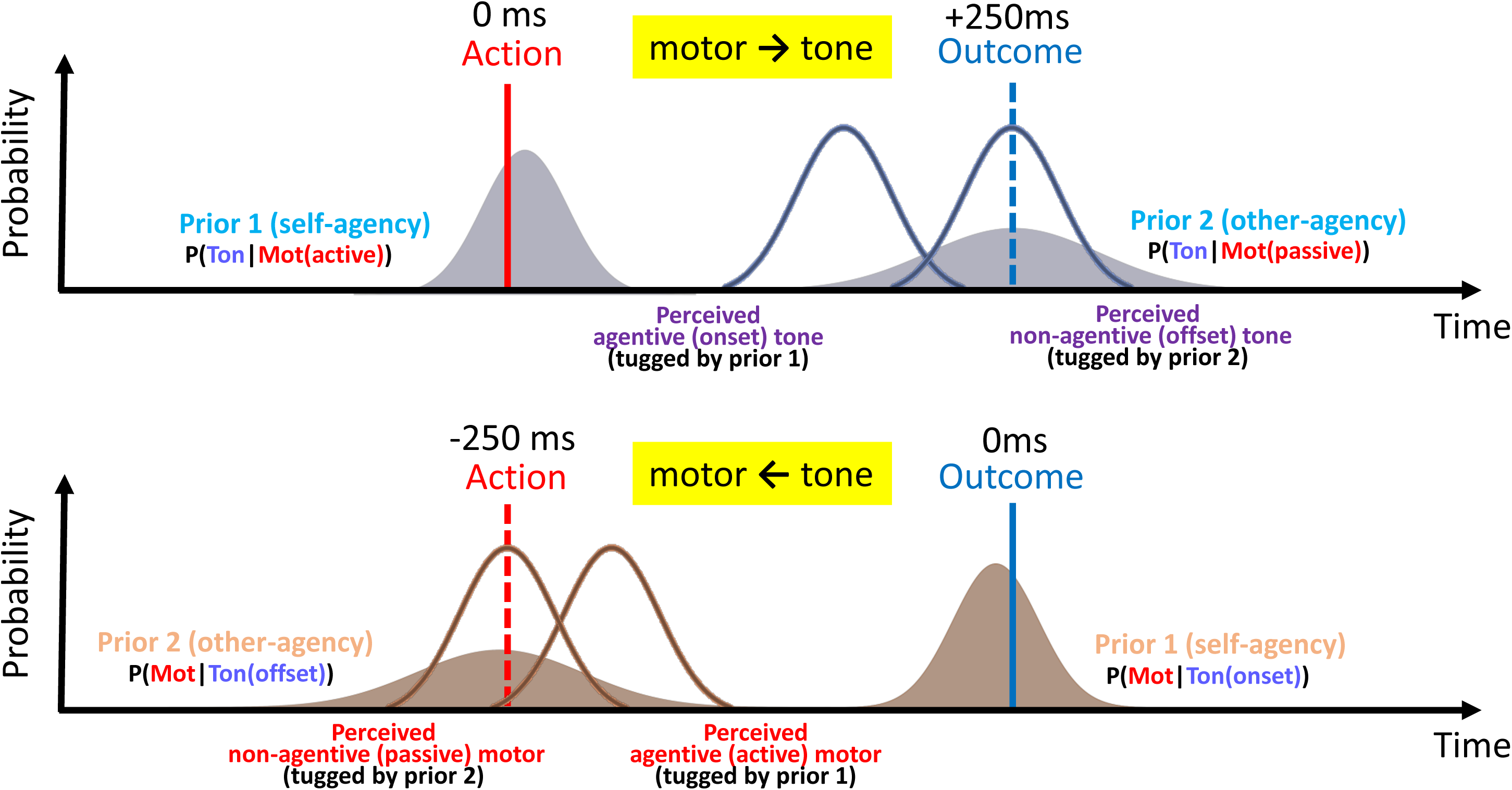
The schematic of Bayesian integration with regard to intentional binding. Note: The distributions in perceived timing under the “agentive” condition are mutually-entrained on the basis of the delay-0 prior (belief) in self-generation. Agentive outcome (onset) is tugged by self-agency prior, while non-agentive outcome (offset) is tugged by other-agency prior (top). Conversely, agentive action (active) is tugged by self prior, while non-agentive one (passive) is tugged by other prior (bottom).

Figure 4 illustrates this coupled uncertainty between action and outcome under the agentive action condition (B: active motor action and onset outcome), compared with presumably uncoupled uncertainty under the baseline condition (A: active motor action and onset tone were presented separately). The significant correlation in SD even under the baseline (*r* = 0.51, *p* < 0.05) suggest general individual differences in sensory uncertainty, indicating that a participant whose reports are more varied in the baseline active motor action condition would also exhibit more varied reports in the baseline onset tone condition (A). However, when these two events were presented at the same time, with delay (B), each reliability was tightly associated with the other (*r* = 0.82, *p* < 0.01), even though each was measured respectively, like they were in (A). When the weighted subtraction (pseudo-distance) from baseline distribution to experimental distribution was calculated as a Kullback-Leibler divergence (e.g., Ais et al., 2016), a shifted peak was observed. Specifically, the timing in active motor action was positively shifted, while that in onset tone was negatively shifted. This was especially true in agentive action-outcome condition (the full KLd matrix is provided in Figure S4). This mutual coupling is theorized in Figure 5, where two categorical priors are assumed. Specifically, that self-agency and other-agency priors will predict the sensory outcome (effect) and the motoric action (cause). Active action predicts self-agency, which further predicts onset tone. Onset tone, in turn, predicts self-agency, which inversely predicts active action. The simple probabilistic inference between cause and effect can be further extended to involve the inference of the “alternative causal structure” or “causal attribution” (Meder, Mayrhofer, & Waldmann, 2014) (e.g., self-other discrimination). Here, the self is learned to be the active generator, as a prior. However, this is not always the case for patients with schizophrenia, e.g., (Haggard, Martin, Taylor-Clarke, Jeannerod, & Franck, 2003; Powers, Mathys, & Corlett, 2017).

#### The coupled (i.e., shared) uncertainty between action and outcome

In order to highlight the coupled uncertainty that was observed only under the agentive action-outcome condition, hierarchical clustering was applied (e.g., Ais et al., 2016) to the distance matrix as characterized by Pearson’s correlation coefficients, where we can assume hierarchical or factorial modulation among experimental conditions (Figure 6). Regarding the mean (left of Figure 6), a clear contrast between tone task and motor task was observed, since the direction of shift in mean should be opposing. Within each the active and passive motor conditions, a unique cluster was observed, including onset/offset conditions. This suggests that active/passive modulation outperforms onset/offset modulation when it comes to the shift of the mean (though this may be due to a block effect). This effect is highly reasonable according to previous studies (i.e., the “intentional” binding effect, (Haggard, 2017) c. f., “causal” binding (Buehner, 2012; Buehner & Humphreys, 2009)). Conversely, when examining the SD (right of Figure 6), we might see a similar clustering pattern except for in the active motor action and onset outcome conditions. This coupling (i.e., *r* = .82) was the only local association across motor and tone tasks (see, for example, the association between active motor action and *offset* outcome condition for comparison, *r* = .50).

**Figure 6.**
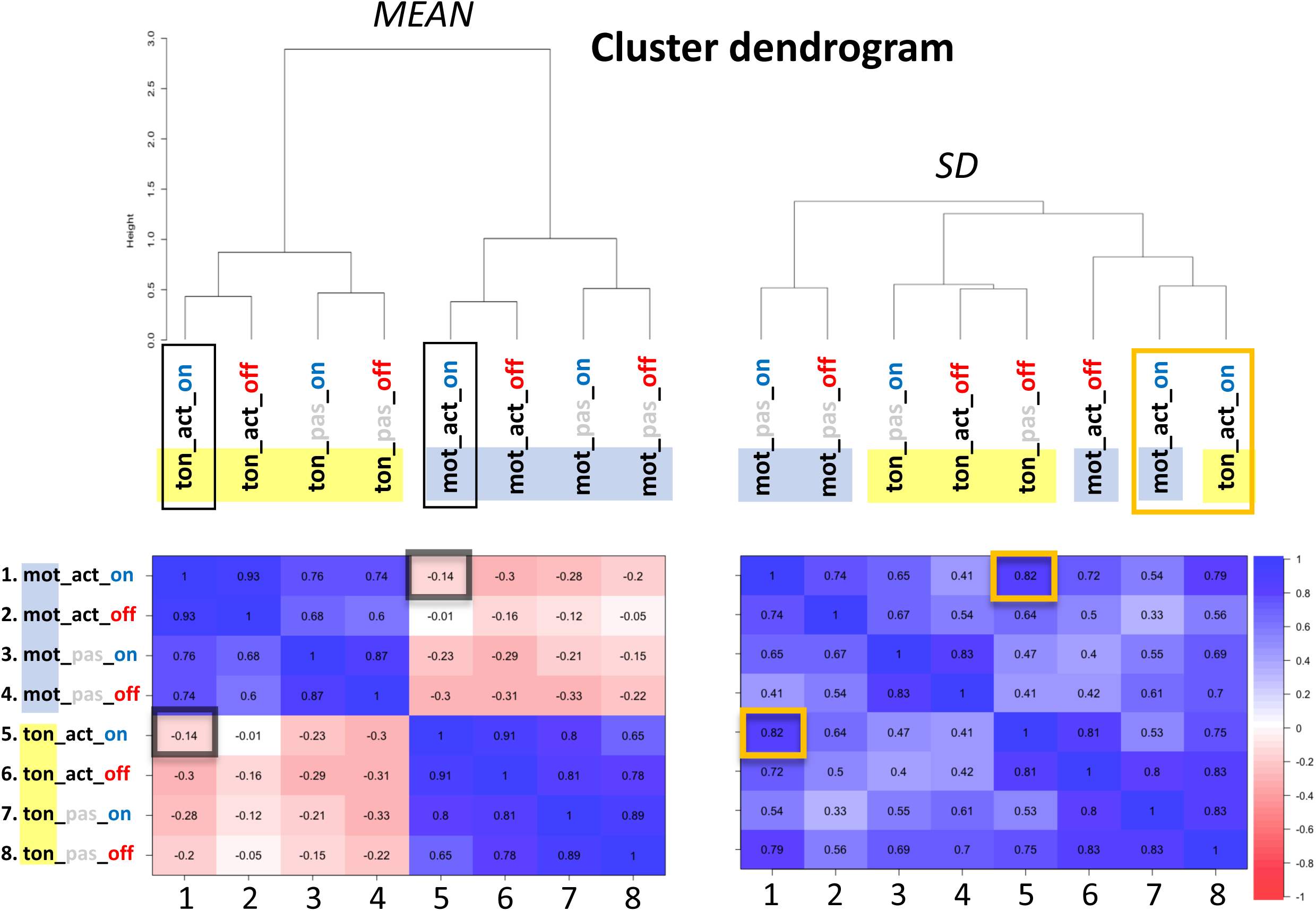
Inter-subject relationships among experimental conditions in means and SDs. Note: Hierarchical clustering with Ward’s method (top) based on Pearson’s correlation (bottom) in means (left) and SDs (right) under each experimental condition.

This suggests that the agentive condition, which requires active production and onset of outcome as an active generator, shares temporal reliability between action and outcome. To illustrate this “network” among all conditions as a whole, the fitted Bayesian causal model (Glymour, 2003) was calculated (Figure 7). Here, the statistical dependency among variables in the whole model are visualized, where the structure, including dependencies and conditional independencies, was determined in an unsupervised manner (i.e., the learner does not distinguish between the dependent and independent variables in the data). This network was characterized similarly as a cluster dendrogram by the difference in tasks, active/passive action, and onset/offset outcome. However, we might see an almost “unicursal” pattern of dependencies in SDs (as a directed acyclic graph is learned in definition), but not in means. Specifically, the coupled (or causal) relationship in SDs between action and outcome under the active-onset condition was again observed. Although convincing interpretations of these kind of models are generally difficult to create (i.e., AI or machine learning approach as “black box”, see (Yarkoni & Westfall, 2017) for discussion), we can at least see the factorial structure in the SDs, compared with the means. This multivariate, mutual relationship is difficult to be *extracted* by traditional statistics (e.g., univariate ANOVA or a correlation for comparison), making it helpful to simulate the results of experimental manipulation for further studies (“interventionism”). This visualization also sheds new light on our action-quantified time perception in terms of sensory precision.

**Figure 7.**
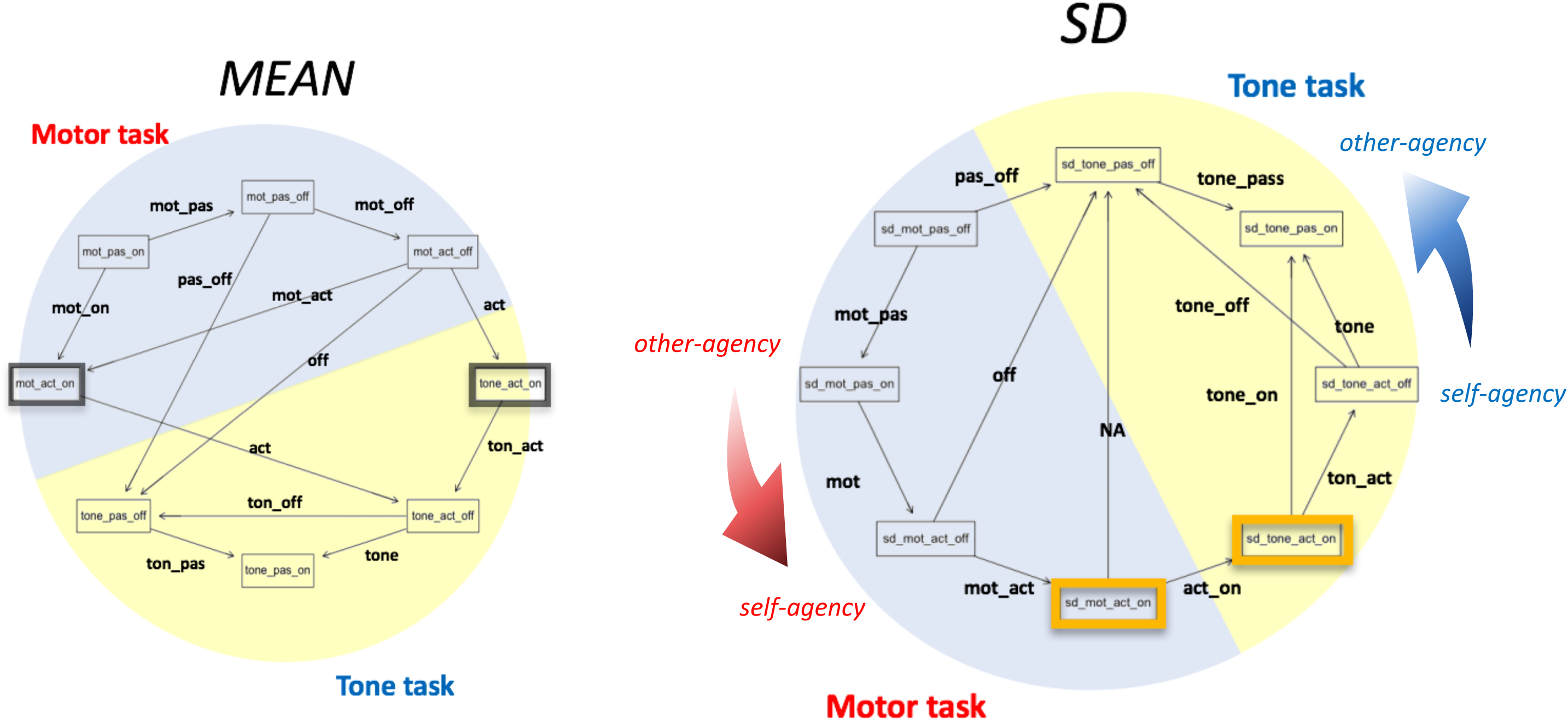
The Bayesian network in means and SDs in Experiment 3. Note: The Bayesian probability relationships among variables were identified by a tabu search algorithm where the arrow indicates “causal” relationship between paired nodes. The text on the edge indicates the shared condition between nodes.

## General Discussion

Results of the study suggest that “agentive action” modulates time perception, both for action itself and for perceptual outcome, where our sensory precision (SD in the current experiments), as well as the detection itself (mean in the current experiments) is somehow altered. Previous studies have suggested that this feeling of agency in action includes *volition* (e.g., active production (Haggard et al., 2002)), *causality* (e.g., delay between action and outcome (Buehner & Humphreys, 2009)), *modality* (e.g., within/across congruency (Engbert et al., 2008; Mifsud & Whitford, 2017)), *context/meaning* of an outcome (e.g., positive/negative emotional event (Yoshie & Haggard, 2013)), and *responsibility* (e.g., obeying social norms (Caspar, Christensen, et al., 2016)). These perceptual and cognitive factors are largely covariate in experimental situations, making it necessary to identify a genuine or minimal factor for agentive action. The current study rolled back to the original and literal meaning of the sense of agency as being as independent agent in the environment. This essential view of agency sets a simple hypothesis that agentive action should apply only to generation, since termination implies another agent in the environment who had already produced the event. The overall results of the current study replicated factors that were previously suggested (*causality* in Experiments 1 and 2, *volition/responsibility* in Experiment 3, *modality* in Experiment 1B, and *context/meaning* in Experiments 2A and 2B) in terms of onset (generation)/offset (termination) agency manipulation. This onset/offset contrast was parameter-free (Experiment 2C) and also task-free (Experiments 1C and 3). In this sense, agentive action requires the generation of a perceptual outcome as well as teleological production. Now we can discuss what happens in our probable (i.e., subjective) sensorium during agentive action.

### Perceptual “event” shifts (Model 1)

We might intuitively interpret the binding effect as the perceived event shifting itself toward the agentive action, independent from the background time flow (e.g., Moore & Obhi, 2012). As a result, we cannot detect delay (e.g., Experiment 1A), we feel a shorter action-outcome interval (Experiment 1C), we feel more agency (e.g., Experiments 1A and 1C), and the exact reports of tone timing are negatively biased (Experiment 3). This model (see Figure 8), however, requires some assumptions. First, we should have a unified and rigid representation of the event or *figure* as a movable perceptual unit (c.f., Model 2). Second, the perceived time flow as *ground* should always be stable (c.f., Model 3). Finally, our precision in time perception should not be modulated by the volitional action (cf., Model 4). Some recent studies have suggested some results that contradict each assumption. We would like to update this simple Model 1 according to the current results, as well as contrasting ideas presented in previous studies.

**Figure 8.**
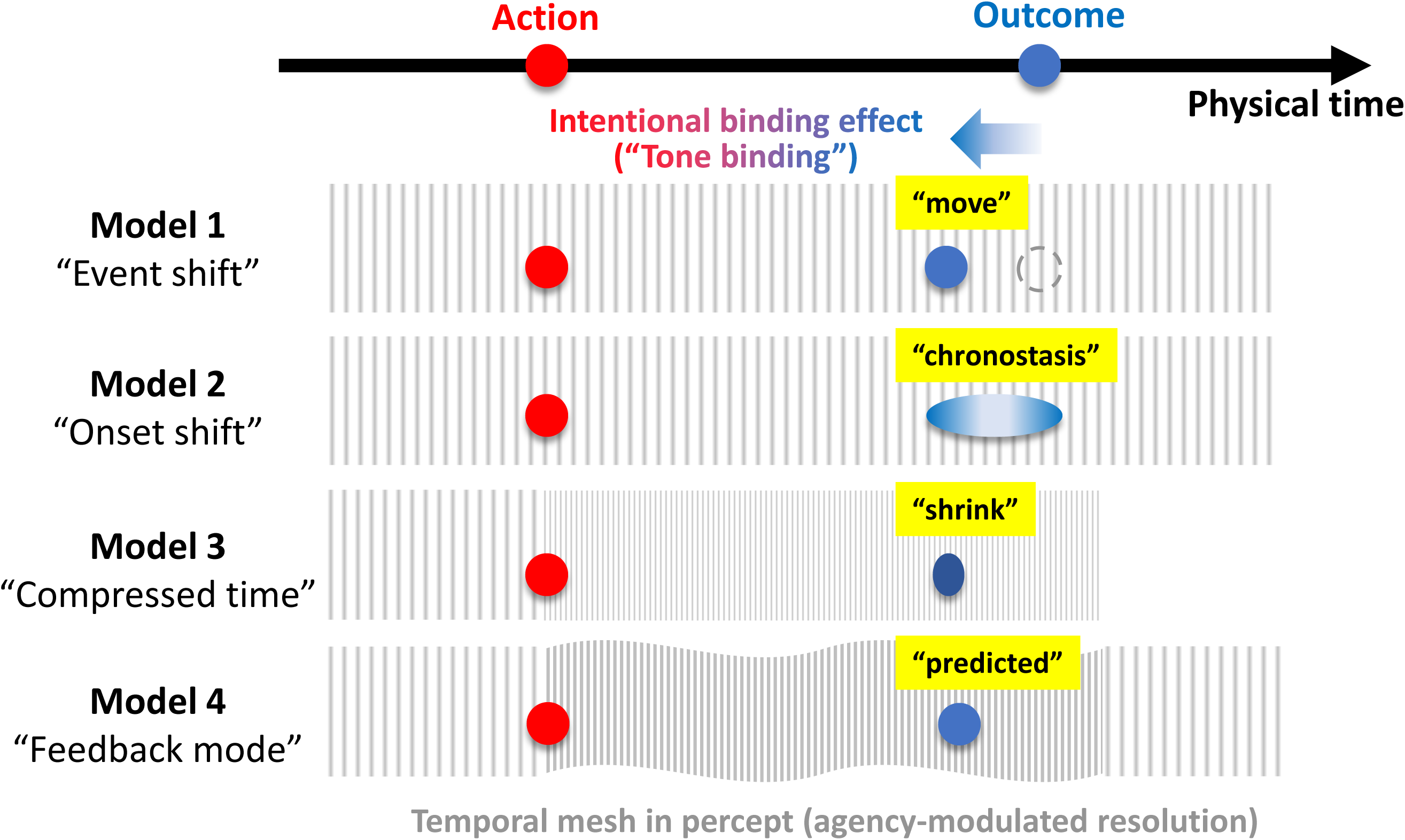
The schematic models of the tone binding, in terms of time. Model 1: Intention attracts the “event.” Model 2: Intention attracts “onset,” but leaves offset. Model 3: Intention attracts “time flow.” Model 4: Intention predicts “agentive outcome” to be detected.

### Only “onset” shifts (Model 2)

A volitional action might attract only the onset timing of a perceptual event, and not the whole event (Model 2 in Figure 8). In this hypothesis, individuals should perceive, not only an onset shift, but also a lengthened duration of the event (Kanai & Watanabe, 2006). One may be reminded of the chronostasis effect, whereby perceptual duration of a visual event following saccadic eye movement is likely to be perceived as longer, possibly due to corollary discharge and a byproduct of saccadic suppression (Yarrow, Haggard, Heal, Brown, & Rothwell, 2001). Analogously to chronostasis, voluntary manual action can lead to the subjective expansion of temporal duration following visual events, but only when the stimulus is presented immediately after the action (Park, Schlag-Rey, & Schlag, 2003). This suggests a possibility that onset timing, but not offset timing, shifts perception toward the action, and as a result, the event duration is expanded. A recent study (Makwana & Srinivasan, 2017) has replicated Park et al.’s finding by extensive experiments, further claiming this only-onset model.

If this model is true, where the perceptual duration of the event dilates, we can further speculate that the perception of onset should be difficult or imprecise, since the actual timing to be judged presents itself without a clear perceptual *edge* (i.e., for a subjectively expanded event). Indeed, the data from Experiment 3 supports this, the variance of reported timing of the tone was significantly larger than baseline, regardless of agency conditions (Figure S5), suggesting that the time perception was less precise in action-outcome situations, especially for tone. This finding, at least partially, is congruent with Model 2. In our mental representation for time, we may not experience a single perceptual event as a consequence of voluntary action.

### Compressed “time flow” (Model 3)

Another possibility is the altered mental time flow both for *figure-ground* (Model 3, Figure 8) where time itself is assumed to be unstable or plastic (i.e., the concept that there is no physical or absolute time proposed in Einstein’s theory of spatio-temporal relativity). We can interpret the binding effect as a compression of the subjective temporal interval between action and outcome, which many studies have demonstrated by using interval estimation (Caspar, Christensen, et al., 2016; Caspar et al., 2015; Engbert et al., 2008; Engbert et al., 2007). In line with this, the results of Experiment 1C indicated that the subjective temporal interval was perceived as shorter in an agentive situation (i.e. tone generation). In addition, Experiment 3 indicated the estimated interval (as composite binding) was compressed only in the most agentive condition (i.e., active-onset). These results may support a compression of mental time flow, but only during agentive action. A widely accepted model for time perception also indicates a pacing rate of neural signals in the “internal clock” (Treisman, 2013). This clock is modulated by several psychological and motoric factors, so that the subjective duration of a stimulus (temporal perceptual resolution) is plastic. Accordingly, a decrease of pacing rate compresses subjective duration, since fewer temporal samples (e.g., neural signals) accumulate within a certain interval. In this sense, intentional binding might be a result of the deceleration of the internal clock, which would coincide with a decrease of perceptual sensitivity or discriminability in time, which is supported by the evidence.

A previous study has indeed suggested that the internal clock dynamically varies its pacing rate triggered by voluntary action within an action-outcome interval (Wenke & Haggard, 2009). In this study, the experimental task was a typical intentional binding procedure (i.e., a tone follows a keypress), but a successive tactile stimulation was introduced onto the participants’ hand immediately after the keypress, at a delayed interval after the keypress, or after the subsequent tone. This study replicated the intentional binding effect in an active-movement condition. Importantly, the tactile temporal discriminability in the active condition was impaired immediately after the keypress, while no such effect was observed later in the keypress-tone interval. Although this dynamically-modulated internal clock may be a neural candidate responsible for the binding effect, care should be taken in reaching this conclusion due to the mixed results reported (Fereday & Buehner, 2017). Taking all of this information into account, an additional model is necessary to tie all of these disparate models together.

### Bayesian “feedback mode” (Model 4)

The above-mentioned models appear to be partially valid. However, previous and the current findings cannot determine which of these models explains the phenomenon best. While future studies will likely examine this issue by employing intentional binding paradigms, duration estimation, and concurrent perceptual sensitivity measures, another updated theoretical model may be helpful for understanding the binding effect, namely, Bayesian integration (i.e., predictive coding) (Adams, Stephan, et al., 2013; Apps & Tsakiris, 2014; Baker et al., 2009; Fletcher & Frith, 2009; Izawa, Asai, & Imamizu, 2016; Lalanne & Lorenceau, 2004; Moreno-Bote, Knill, & Pouget, 2011; Moutoussis et al., 2014). Agency has already been discussed in terms of prediction or prediction error (e.g., in simple comparator or optimal cue integration (Synofzik, Vosgerau, & Newen, 2008; Wolpe et al., 2013)). Though this view sounds reasonable, especially for continuous motor tasks, since this framework derives from the internal model of motor control (Wolpert, 1997; Wolpert, Miall, & Kawato, 1998), recent binding studies also refer to prediction (Moore & Haggard, 2008; Yoshie & Haggard, 2017)). Therefore, we can combine a “single action” task and a “continuous motor” task in terms of spatio-temporal prediction (Blakemore et al., 1999) by Bayesian predictive coding (Picard & Friston, 2014; Rao & Ballard, 1999).

We have learned the internal model for predicting sensory outcomes, and further, for controlling our own body. If prediction error is detected, we attribute this error (which is generally small) to ourselves, in order to update the internal model for the self in real-time (e.g., we can modify our own motor performance). Alternatively, we might use this “(often big) surprise” as an engine to better understand others, and the world at large (e.g., we can understand the “law” on an artificial force field (McIntyre, Zago, Berthoz, & Lacquaniti, 2001)). This implies that the attributed source (self or other) is selected/weighted on the basis of relative reliabilities (see Figure 5). Here, we have two options to eliminate the encountered prediction error: updating the prediction so that error is reduced (belief renovation), or recalibration of the sensation (active inference) (Adams, Shipp, & Friston, 2013). Both, in reality, might coincide with being weighted, so that the distribution for prediction and for sensation are mutually skewed. The current study suggests that the self is learned as the teleological generator. In other words, we have a strong prediction or prior where “I” should be the entity that must produce sensation immediately by volitional action (Bays et al., 2005). Therefore, in the case of the current experiments, the prediction errors to be eliminated included delay (under the agentive condition), passive action (against onset outcome), and offset outcome (against active action). While delay can be reduced by active inference (an actual delay is no longer considered a delay), the passive action and offset outcome are believed to be attributed to an agent other than the self, resulting in no need to recalibrate one’s own sensorium.

Essentially, any perception cannot be prediction (or bias) free (K. J. Friston et al., 2014; Shipp et al., 2013). Specifically, in an action-outcome situation, action and perception are tightly coupled in our “sensorimotor” system, where traditional ideomotor theory meets the modern Bayesian integration account (Caspar, Desantis, et al., 2016; Khalighinejad & Haggard, 2016). We might name this model a “feedback mode,” with the ability to include previous models (Model 4 in Figure 8), suggesting that a specific action predicts a specific outcome (and vice versa). Therefore, during agentive action, we have a special conscious state to wait/prepare for the predicted sensory outcome (e.g., return-trip effect (Maglio & Kwok, 2016; van de Ven, van Rijswijk, & Roy, 2011)). The previous models have suggested that our subjective “duration” as a *figure* is dilated (Model 2), but “interval” as *ground* might be compressed (Model 3). This modulation can also dynamically change depending on the phase of bodily movement. Studies have observed a “clock-up” phenomenon (i.e., dilated duration and enhanced perceptual resolution of time) in motor preparation (Hagura, Kanai, Orgs, & Haggard, 2012) and execution periods (Imaizumi & Asai, 2017; Press, Berlot, Bird, Ivry, & Cook, 2014). In contrast, a “clock-down” phenomenon has been observed in post-execution period (Tomassini, Gori, Baud-Bovy, Sandini, & Morrone, 2014; Wenke & Haggard, 2009), although contrary results of these phenomena have also been reported (Fereday & Buehner, 2017). Perceptual inhibition (or enhancement) for the targeted stimuli, known as sensory attenuation, is also still controversial (Eliades & Wang, 2008; Kaiser & Schutz-Bosbach, 2018; Kilteni, Andersson, Houborg, & Ehrsson, 2018; Poulet & Hedwig, 2002; Wen, Brann, Di Costa, & Haggard, 2018). The field still does not know exactly what is happening during agentive action, but a “feedback mode” model suggests that all depend on how priors are selected/learned. It is possible that the phrase “to be precise, the details don’t matter” is applicable when encountering the trade-off between prediction precision and information gain (Kwisthout, Bekkering, & van Rooij, 2017), where the “categorical” probability or inference might be prioritized, as can be seen by the current results (e.g., *self* or *other* in Figure 5). This subjective categorical attribution observed in a bidirectional loop between teleology and generativity should, in turn, affect our perception, including delusions/hallucinations (hierarchical top-down modulation, e.g., Powers et al., 2017). Future study is necessary to visualize how our large of an impact priors have on our perceptions and actions, as the current study focused primarily on perceptual variance as prior-biased sensory reliability/uncertainty.

## Acknowledgements

The authors are grateful to Dr. Patrick Haggard (The University College London) for his insightful comments on the experimental procedure and also on an earlier manuscript. T.A. was supported by JSPS KAKENHI Grant Number 17K13971. S.I. was supported by JSPS KAKENHI Grant Number 16J00411 and 17K12701. H.I. were supported by JSPS KAKENHI Grant Numbers 26120002 and 18H01098. The authors are also grateful to Yu Miyawaki and Wataru Miyakawa for their partial support in data collection. The research results have been achieved by “Research and development of technology for enhancing functional recovery of elderly and disabled people based on non-invasive brain imaging and robotic assistive devices”, the Commissioned Research of National Institute of Information and Communications Technology (NICT), JAPAN and also Japan Agency for Medical Research and Development (AMED) under Grant Number JP18dm0307008.

## Author contribution statement

T.A., S.I. and H.I. designed the study; T.A. collected its data, and analyzed the data; T.A., S.I. and H.I. wrote the manuscript, reviewed, and approved the final version of the manuscript.

---

**Figure S1.**
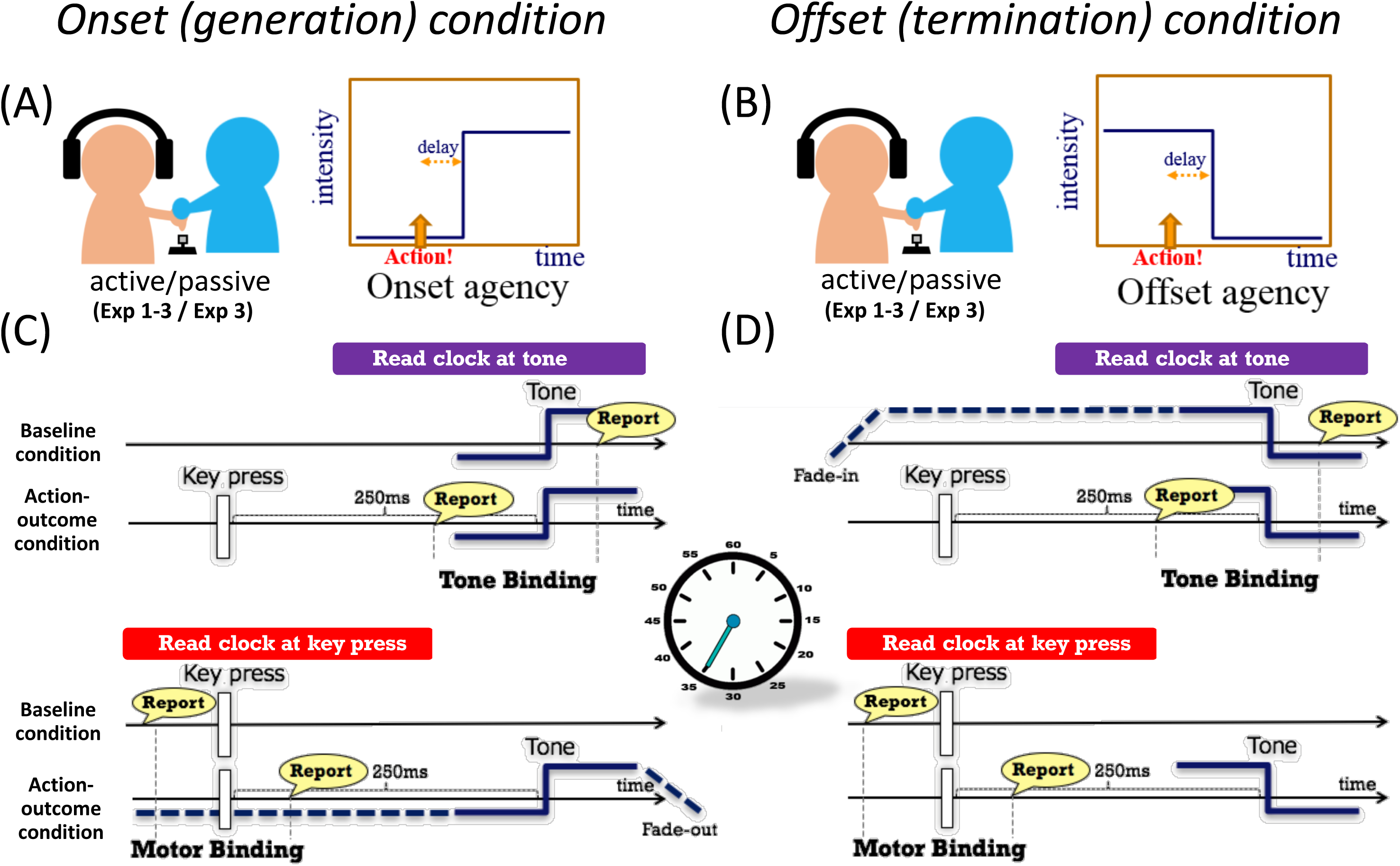
The experimental manipulation of on/offset agency in terms of delay. Note: Agency judgement or delay detection was required in Experiments 1 and2 (A, B), while a reading-clock task was administered in Experiment 3 (C, D) in socially active/passive x onset/offset situations (left or right panels).

**Figure S2.**
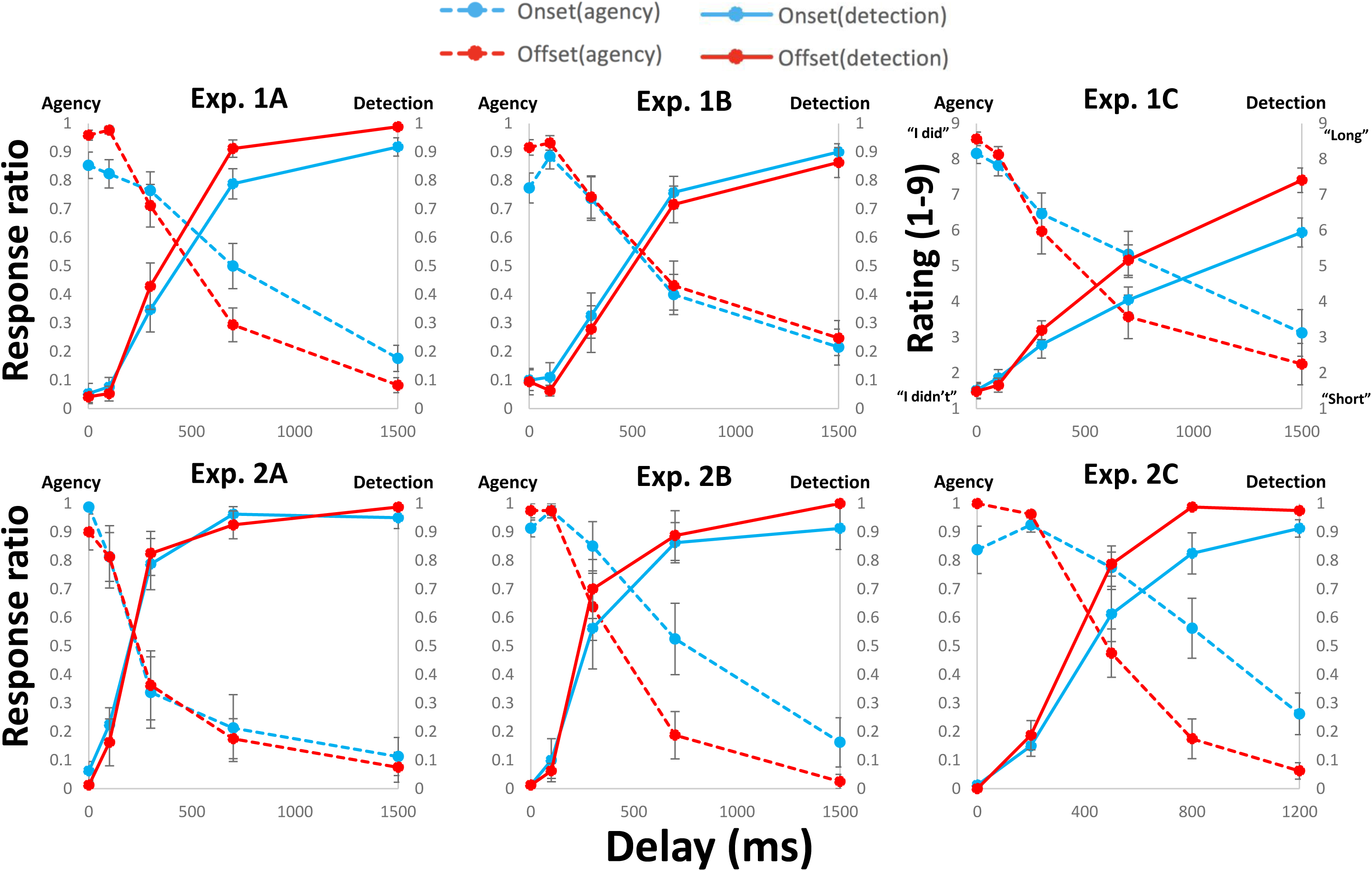
The mean participants’ raw reports in Experiments 1 and 2. Note: Error bars indicate ±1 SE.

**Figure S3.**
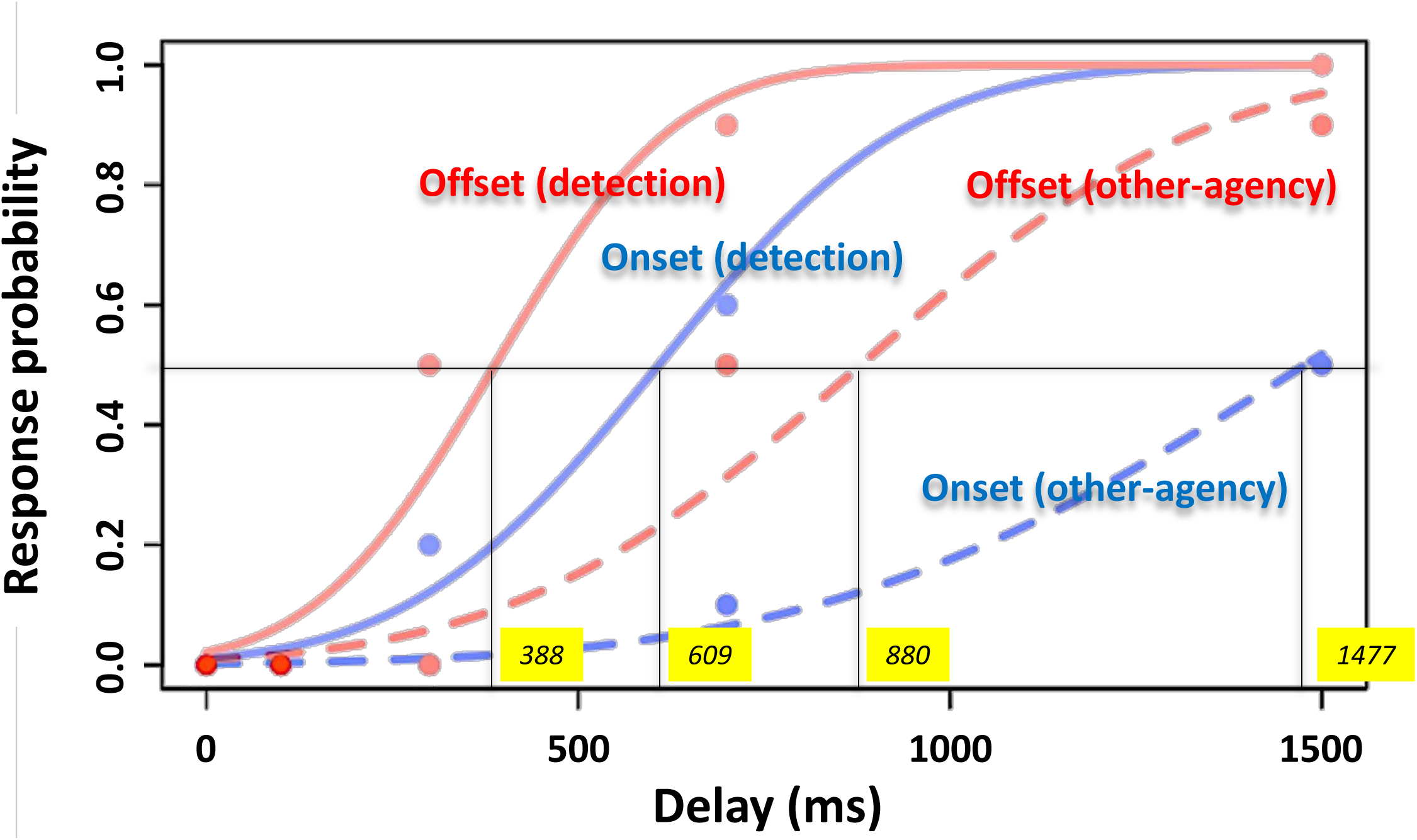
The fitted psychometric functions and estimated 50% thresholds. Note: The result of a typical participant was plotted. The probit regression (cumulative normal distribution) was applied to each participant’s raw binary reports. 50% thresholds were estimated individually (yellow).

**Figure S4.**
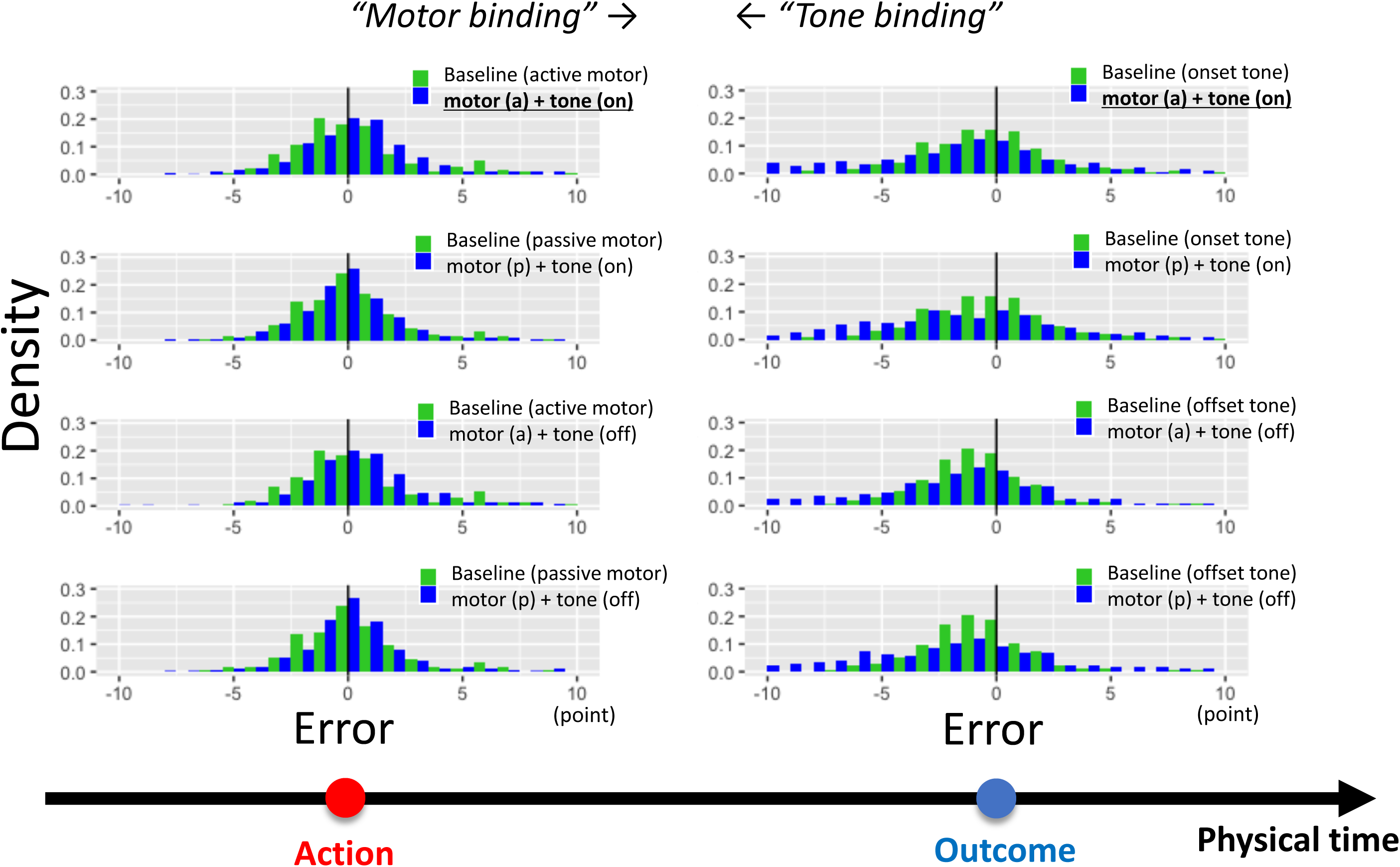
The grand raw reports on a dodged histogram for all 12 conditions in Experiment 3. Note: Each baseline condition (4 conditions) are plotted twice for the comparison with experimental conditions. <u>Underbars</u> represent the experimental condition of interest (see main text for details).

**Figure S5.**
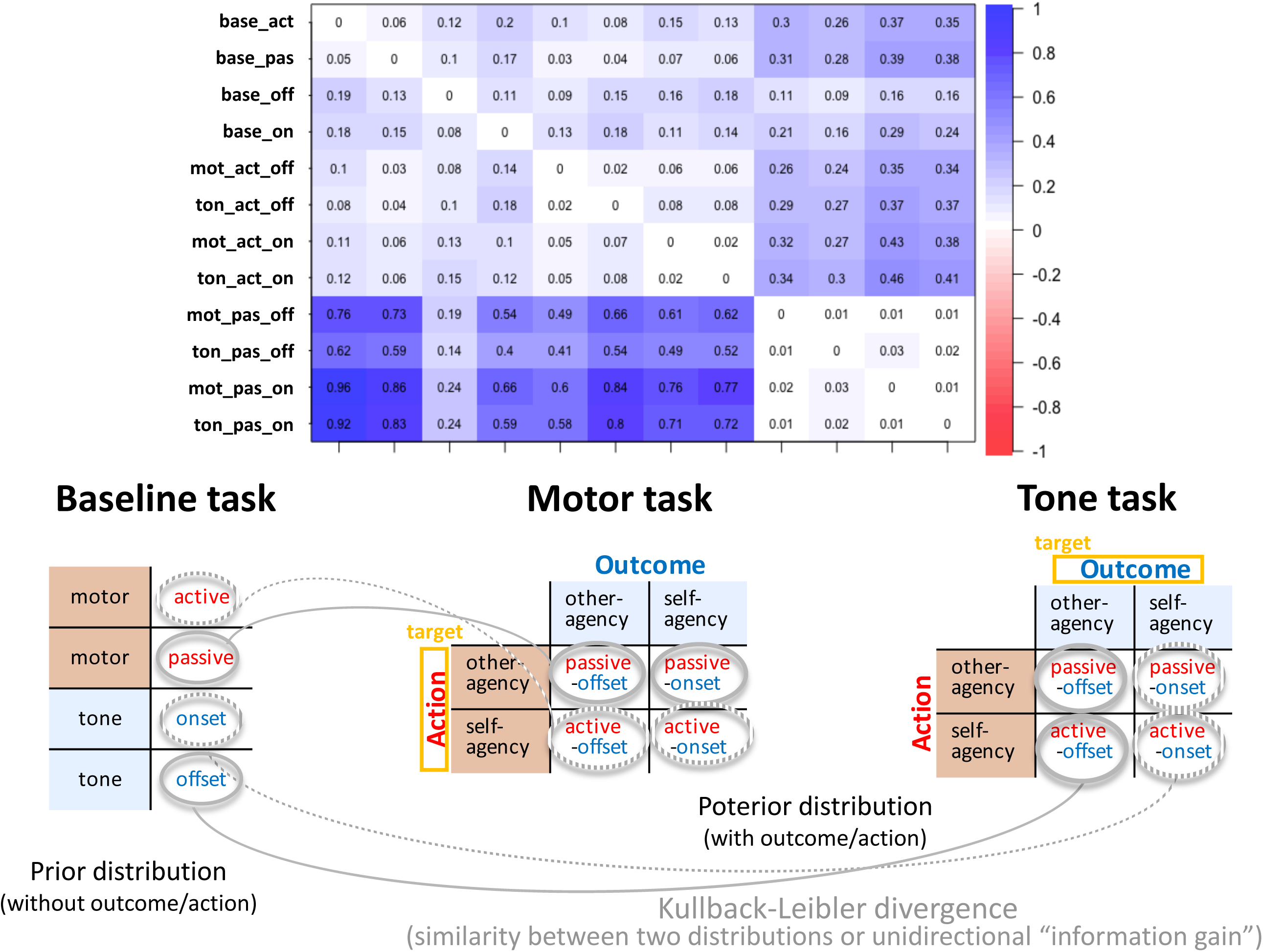
The summed KLd from baseline as prior, to experimental task as posterior distribution. Note: The KLd is the prior-weighted pseudo-distance between two probability distributions at 1000 points within the scoring range. The original discrete relative frequency (see Figure S3) was fitted by a density estimation with a Gaussian kernel window. The summed KLd (top) should be a positive value in definition, though raw negative values are also observable, but only at some points (see Figure 4EF).

**Figure S6.**
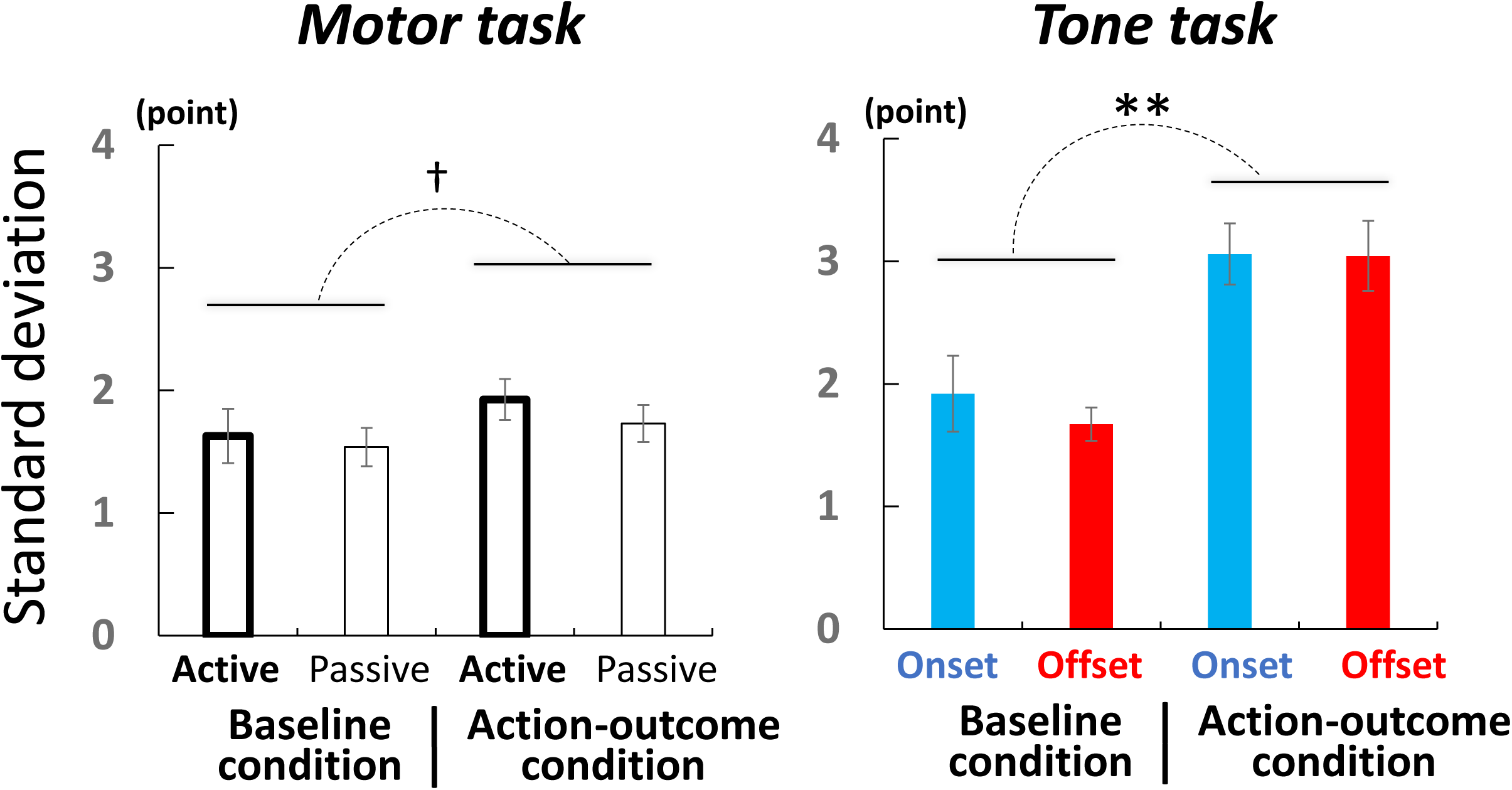
Participants’ mean SD in each condition of Experiment 3. Note: Under the action-outcome condition in the motor task, active or passive condition combines onset/offset conditions for comparison with baseline, while under action-outcome condition in tone task, onset/offset condition combines active and passive conditions as well. Error bars represents ± 1 SE. ** p < .01, * p < .05, † p < .10.

**Figure S7.**
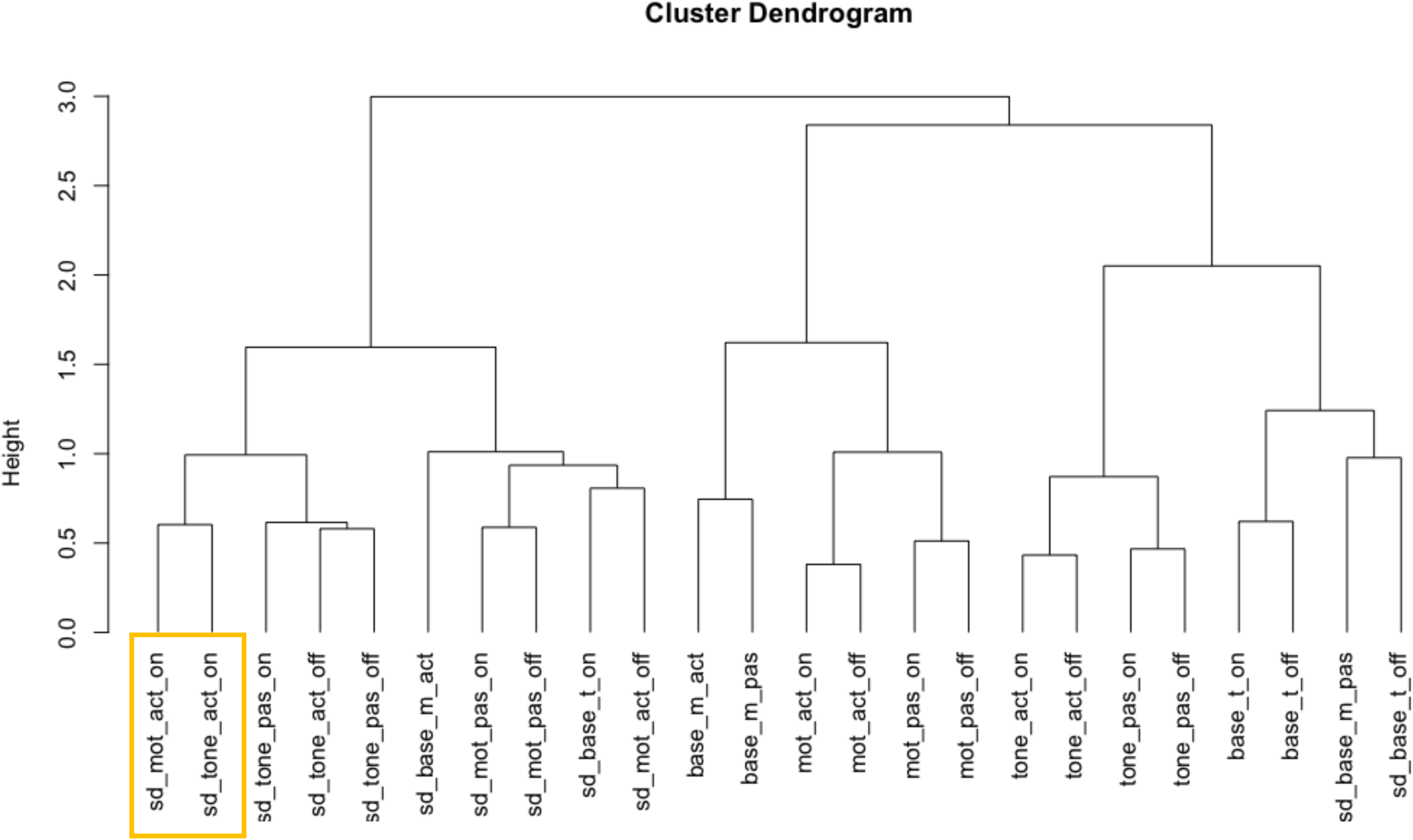
Full cluster dendrogram across means and SDs in Experiment 3.

**Figure S8.**
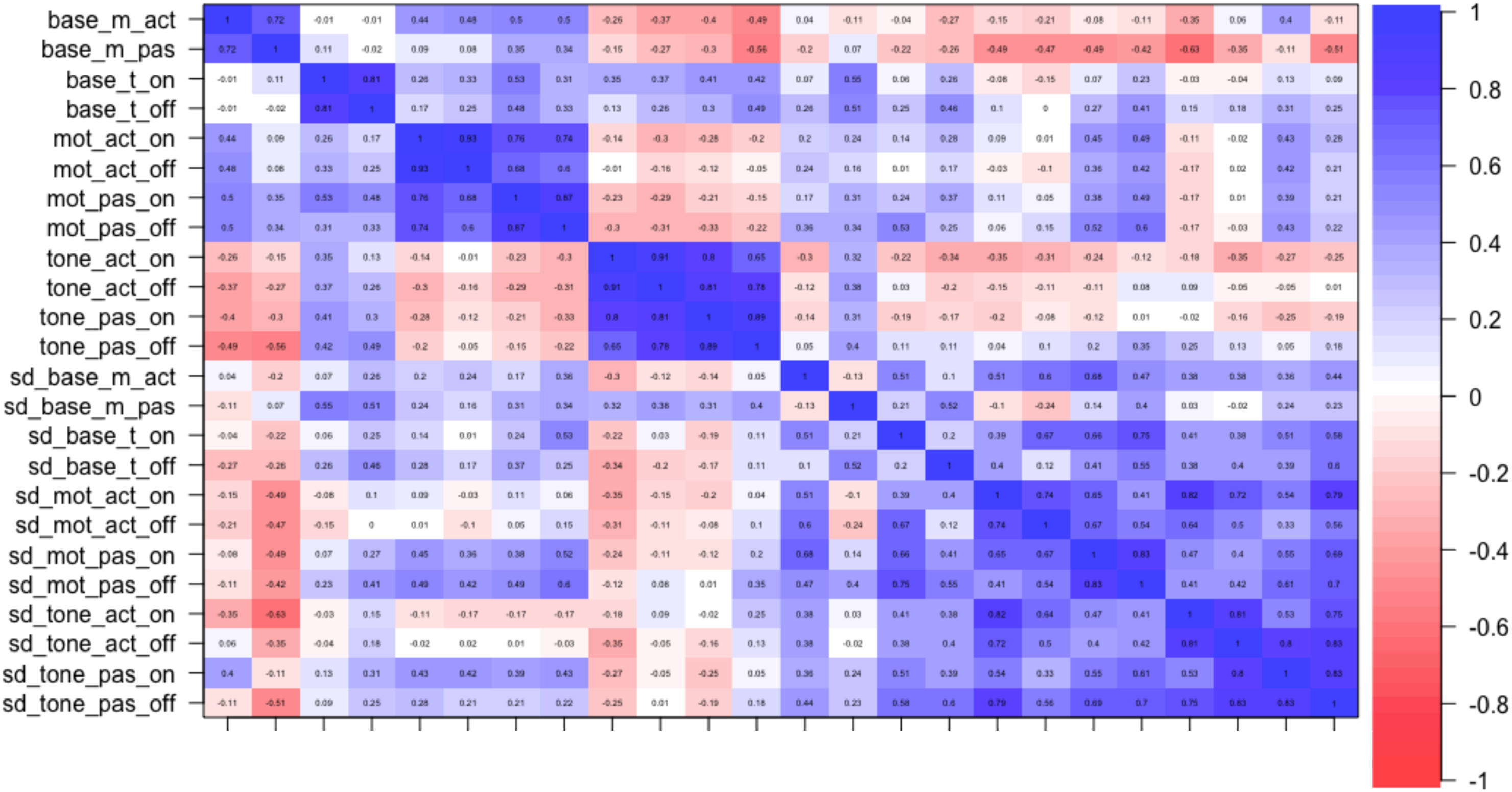
The full correlation matrix across meanss and SDs in Experiment 3.

**Table S1.**
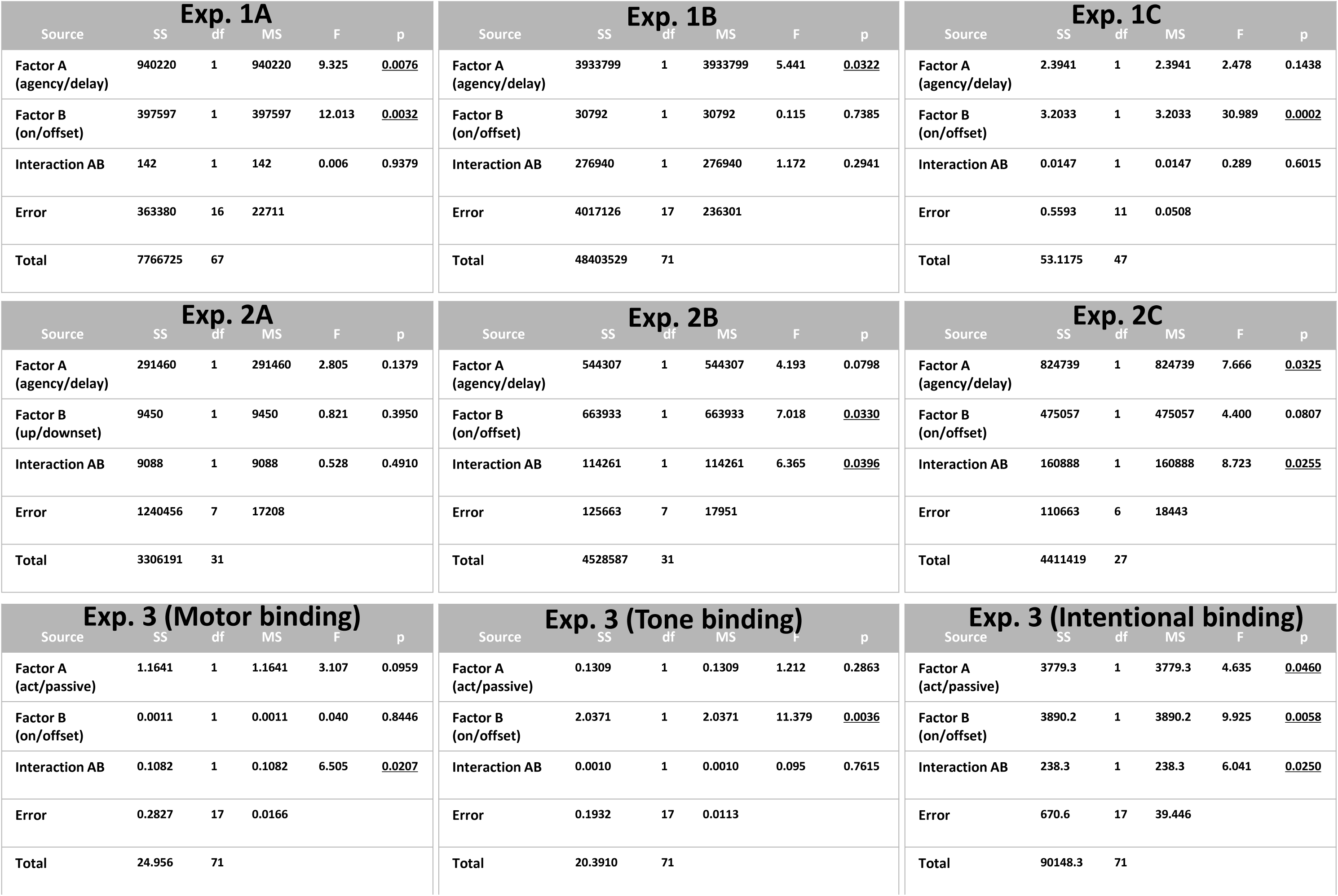
Full ANOVA results of Experiments 1–3. Note: Underbars represent the statistical significance (p < .05).

**Table S2.** The means and SDs of reports in each condition in Experiment 3.

| condition | MEAN |  | SD |  |
| --- | --- | --- | --- | --- |
|  | mean | SE | mean | SE |
| base_mot_act | 0.009 | 0.129 | 1.629 | 0.221 |
| base_mot_pas | 0.172 | 0.135 | 1.538 | 0.157 |
| base_ton_on | 0.142 | 0.131 | 1.919 | 0.31 |
| base_ton_off | -0.22 | 0.116 | 1.672 | 0.135 |
| mot_act_on | 0.211 | 0.074 | 2.033 | 0.191 |
| mot_act_off | 0.126 | 0.1 | 1.82 | 0.17 |
| mot_pas_on | 0.042 | 0.104 | 1.72 | 0.16 |
| mot_pas_off | 0.111 | 0.086 | 1.739 | 0.156 |
| tone_act_on | -0.25 | 0.11 | 2.962 | 0.279 |
| tone_act_off | -0.27 | 0.108 | 2.983 | 0.275 |
| tone_pas_on | -0.15 | 0.112 | 3.156 | 0.292 |
| tone_pas_off | -0.19 | 0.121 | 3.105 | 0.323 |

(point)

## References

Corlett, P. I Predict, Therefore I Am: Perturbed Predictive Coding Under Ketamine and in Schizophrenia. Biological Psychiatry, 81, 465–466 (2017).

Frith, C. D., Haggard, P. Volition and the Brain - Revisiting a Classic Experimental Study. Trends Neurosci. 41, 405–407 (2018).

Gallagher, S. Philosophical conceptions of the self: implications for cognitive science. Trends Cogn. Sci. 4, 14–21 (2000).

Haggard, P. The Neurocognitive Bases of Human Volition. Annu Rev Psychol. (in press)

## References

Adams, R. A., Shipp, S., & Friston, K. J. (2013). Predictions not commands: active inference in the motor system. Brain Structure and Function, 218(3), 611–643. doi:10.1007/s00429-012-0475-5

Adams, R. A., Stephan, K. E., Brown, H. R., Frith, C. D., & Friston, K. J. (2013). The computational anatomy of psychosis. Frontiers in Psychiatry, 4, 47. doi:10.3389/fpsyt.2013.00047

Ais, J., Zylberberg, A., Barttfeld, P., & Sigman, M. (2016). Individual consistency in the accuracy and distribution of confidence judgments. Cognition, 146, 377–386. doi:10.1016/j.cognition.2015.10.006

Apps, M. A., & Tsakiris, M. (2014). The free-energy self: a predictive coding account of self-recognition. Neuroscience and Biobehavioral Reviews, 41, 85–97. doi:10.1016/j.neubiorev.2013.01.029

Asai, T., & Tanno, Y. (2007). The relationship between the sense of self-agency and schizotypal personality traits. Journal of Motor Behavior, 39(3), 162–168. doi:10.3200/JMBR.39.3.162-168

Asai, T., & Tanno, Y. (2008). Highly schizotypal students have a weaker sense of self-agency. Psychiatry and Clinical Neurosciences, 62(1), 115–119. doi:10.1111/j.1440-1819.2007.01768.x

Asai, T., & Tanno, Y. (2013). Why must we attribute our own action to ourselves? Auditory hallucination like-experiences as the results both from the explicit self-other attribution and implicit regulation in speech. Psychiatry Research, 207(3), 179–188. doi:10.1016/j.psychres.2012.09.055

Baker, C. L., Saxe, R., & Tenenbaum, J. B. (2009). Action understanding as inverse planning. Cognition, 113(3), 329–349. doi:10.1016/j.cognition.2009.07.005

Bays, P. M., Wolpert, D. M., & Flanagan, J. R. (2005). Perception of the consequences of self-action is temporally tuned and event driven. Current Biology, 15(12), 1125–1128. doi:10.1016/j.cub.2005.05.023

Beck, B., Di Costa, S., & Haggard, P. (2017). Having control over the external world increases the implicit sense of agency. Cognition, 162, 54–60. doi:10.1016/j.cognition.2017.02.002

Beyer, F., Sidarus, N., Bonicalzi, S., & Haggard, P. (2017). Beyond self-serving bias: diffusion of responsibility reduces sense of agency and outcome monitoring. Social Cognitive and Affective Neuroscience, 12(1), 138–145. doi:10.1093/scan/nsw160

Beyer, F., Sidarus, N., Fleming, S., & Haggard, P. (2018). Losing control in social situations: How the presence of others affects neural processes related to sense of agency. eNeuro, 5(1). doi:10.1523/ENEURO.0336-17.2018

Blakemore, S. J., Frith, C. D., & Wolpert, D. M. (1999). Spatio-temporal prediction modulates the perception of self-produced stimuli. Journal of Cognitive Neuroscience, 11(5), 551–559. doi:10.1162/089892999563607

Borhani, K., Beck, B., & Haggard, P. (2017). Choosing, doing, and controlling: Implicit sense of agency over somatosensory events. Psychological Science, 28(7), 882–893. doi:10.1177/0956797617697693

Brooks, J. X., Carriot, J., & Cullen, K. E. (2015). Learning to expect the unexpected: rapid updating in primate cerebellum during voluntary self-motion. Nature Neuroscience, 18(9), 1310–1317. doi:10.1038/nn.4077

Brown, H., Adams, R. A., Parees, I., Edwards, M., & Friston, K. (2013). Active inference, sensory attenuation and illusions. Cognitive Processing, 14(4), 411–427. doi:10.1007/s10339-013-0571-3

Buehner, M. J. (2012). Understanding the past, predicting the future: causation, not intentional action, is the root of temporal binding. Psychological Science, 23(12), 1490–1497. doi:10.1177/0956797612444612

Buehner, M. J., & Humphreys, G. R. (2009). Causal binding of actions to their effects. Psychological Science, 20(10), 1221–1228. doi:10.1111/j.1467-9280.2009.02435.x

Caspar, E. A., Christensen, J. F., Cleeremans, A., & Haggard, P. (2016). Coercion changes the sense of agency in the human brain. Current Biology, 26(5), 585–592. doi:10.1016/j.cub.2015.12.067

Caspar, E. A., Cleeremans, A., & Haggard, P. (2015). The relationship between human agency and embodiment. Consciousness and Cognition, 33, 226–236. doi:10.1016/j.concog.2015.01.007

Caspar, E. A., Desantis, A., Dienes, Z., Cleeremans, A., & Haggard, P. (2016). The sense of agency as tracking control. PLoS ONE, 11(10), e0163892. doi:10.1371/journal.pone.0163892

Cavazzana, A., Begliomini, C., & Bisiacchi, P. S. (2017). Intentional binding as a marker of agency across the lifespan. Consciousness and Cognition, 52, 104–114. doi:10.1016/j.concog.2017.04.016

Crivelli, D., & Balconi, M. (2017). The agent brain: A review of non-invasive brain stimulation studies on sensing agency. Frontiers in Behavioral Neuroscience, 11, 229. doi:10.3389/fnbeh.2017.00229

Daprati, E., Franck, N., Georgieff, N., Proust, J., Pacherie, E., Dalery, J., & Jeannerod, M. (1997). Looking for the agent: an investigation into consciousness of action and self-consciousness in schizophrenic patients. Cognition, 65(1), 71–86. doi:10.1016/s0010-0277(97)00039-5

Desantis, A., & Haggard, P. (2016). Action-outcome learning and prediction shape the window of simultaneity of audiovisual outcomes. Cognition, 153, 33–42. doi:10.1016/j.cognition.2016.03.009

Desantis, A., Hughes, G., & Waszak, F. (2012). Intentional binding is driven by the mere presence of an action and not by motor prediction. PLoS ONE, 7(1), e29557. doi:10.1371/journal.pone.0029557

Dewey, J. A., & Knoblich, G. (2014). Do implicit and explicit measures of the sense of agency measure the same thing? PLoS ONE, 9(10), e110118. doi:10.1371/journal.pone.0110118

Ebert, J. P., & Wegner, D. M. (2010). Time warp: authorship shapes the perceived timing of actions and events. Consciousness and Cognition, 19(1), 481–489. doi:10.1016/j.concog.2009.10.002

Eliades, S. J., & Wang, X. (2008). Neural substrates of vocalization feedback monitoring in primate auditory cortex. Nature, 453(7198), 1102–1106. doi:10.1038/nature06910

Engbert, K., Wohlschlager, A., & Haggard, P. (2008). Who is causing what? The sense of agency is relational and efferent-triggered. Cognition, 107(2), 693–704. doi:10.1016/j.cognition.2007.07.021

Engbert, K., Wohlschlager, A., Thomas, R., & Haggard, P. (2007). Agency, subjective time, and other minds. Journal of Experimental Psychology: Human Perception and Performance, 33(6), 1261–1268. doi:10.1037/0096-1523.33.6.1261

Farrer, C., Valentin, G., & Hupe, J. M. (2013). The time windows of the sense of agency. Consciousness and Cognition, 22(4), 1431–1441. doi:10.1016/j.concog.2013.09.010

Fereday, R., & Buehner, M. J. (2017). Temporal binding and internal clocks: No evidence for general pacemaker slowing. Journal of Experimental Psychology: Human Perception and Performance, 43(5), 971–985. doi:10.1037/xhp0000370

Fletcher, P. C., & Frith, C. D. (2009). Perceiving is believing: a Bayesian approach to explaining the positive symptoms of schizophrenia. Nature Reviews: Neuroscience, 10(1), 48–58. doi:10.1038/nrn2536

Friston, K. (2010). Is the free-energy principle neurocentric? Nature Reviews: Neuroscience, 11(8), 605. doi:10.1038/nrn2787-c2

Friston, K. J., Stephan, K. E., Montague, R., & Dolan, R. J. (2014). Computational psychiatry: the brain as a phantastic organ. The Lancet Psychiatry, 1(2), 148–158. doi:10.1016/S2215-0366(14)70275-5

Gallagher, S. (2000). Philosophical conceptions of the self: implications for cognitive science. Trends in Cognitive Sciences, 4(1), 14–21. doi:10.1016/S1364-6613(99)01417-5

Gentsch, A., & Synofzik, M. (2014). Affective coding: the emotional dimension of agency. Frontiers in Human Neuroscience, 8, 608. doi:10.3389/fnhum.2014.00608

Gentsch, A., Weber, A., Synofzik, M., Vosgerau, G., & Schutz-Bosbach, S. (2016). Towards a common framework of grounded action cognition: Relating motor control, perception and cognition. Cognition, 146, 81–89. doi:10.1016/j.cognition.2015.09.010

Glymour, C. (2003). Learning, prediction and causal Bayes nets. Trends in Cognitive Sciences, 7(1), 43–48. doi:10.1016/S1364-6613(02)00009-8

Haggard, P. (2017). Sense of agency in the human brain. Nature Reviews: Neuroscience, 18(4), 196–207. doi:10.1038/nrn.2017.14

Haggard, P., Clark, S., & Kalogeras, J. (2002). Voluntary action and conscious awareness. Nature Neuroscience, 5(4), 382–385. doi:10.1038/nn827

Haggard, P., Martin, F., Taylor-Clarke, M., Jeannerod, M., & Franck, N. (2003). Awareness of action in schizophrenia. Neuroreport, 14(7), 1081–1085. doi:10.1097/01.wnr.0000073684.00308.c0

Hagura, N., Kanai, R., Orgs, G., & Haggard, P. (2012). Ready steady slow: action preparation slows the subjective passage of time. Proceedings: Biological Sciences, 279(1746), 4399–4406. doi:10.1098/rspb.2012.1339

Hansen, K. D., Gentry, J., Long, L., Gentleman, R., Falcon, S., Hahne, F., & Sarkar, D. (2017). Rgraphviz: Provides plotting capabilities for R graph objects. R package version 2.22.0. Retrieved from http://bioconductor.org/packages/Rgraphviz/

Hon, N., Seow, Y. Y., & Pereira, D. (2018). Outside influence: The sense of agency takes into account what is in our surroundings. Acta Psychologica. doi:10.1016/j.actpsy.2018.03.004

Hove, M. J., Fairhurst, M. T., Kotz, S. A., & Keller, P. E. (2013). Synchronizing with auditory and visual rhythms: an fMRI assessment of modality differences and modality appropriateness. NeuroImage, 67, 313–321. doi:10.1016/j.neuroimage.2012.11.032

Hubel, D. H., & Wiesel, T. N. (1962). Receptive fields, binocular interaction and functional architecture in the cat’s visual cortex. Journal of Physiology, 160, 106–154. doi:10.1113/jphysiol.1962.sp006837

Imaizumi, S., & Asai, T. (2017). My action lasts longer: Potential link between subjective time and agency during voluntary action. Consciousness and Cognition, 51, 243–257. doi:10.1016/j.concog.2017.04.006

Izawa, J., Asai, T., & Imamizu, H. (2016). Computational motor control as a window to understanding schizophrenia. Neuroscience Research, 104, 44–51. doi:10.1016/j.neures.2015.11.004

Jancke, L., Loose, R., Lutz, K., Specht, K., & Shah, N. J. (2000). Cortical activations during paced finger-tapping applying visual and auditory pacing stimuli. Cognitive Brain Research, 10(1-2), 51–66. doi:10.1016/S0926-6410(00)00022-7

Jeannerod, M. (2003). The mechanism of self-recognition in humans. Behavioural Brain Research, 142(1-2), 1–15. doi:10.1016/S0166-4328(02)00384-4

Kaiser, J., & Schutz-Bosbach, S. (2018). Sensory attenuation of self-produced signals does not rely on self-specific motor predictions. European Journal of Neuroscience. doi:10.1111/ejn.13931

Kanai, R., & Watanabe, M. (2006). Visual onset expands subjective time. Perception and Psychophysics, 68(7), 1113–1123. doi:10.3758/BF03193714

Khalighinejad, N., & Haggard, P. (2016). Extending experiences of voluntary action by association. Proceedings of the National Academy of Sciences of the United States of America, 113(31), 8867–8872. doi:10.1073/pnas.1521223113

Khalighinejad, N., Schurger, A., Desantis, A., Zmigrod, L., & Haggard, P. (2018). Precursor processes of human self-initiated action. NeuroImage, 165, 35–47. doi:10.1016/j.neuroimage.2017.09.057

Kilteni, K., Andersson, B. J., Houborg, C., & Ehrsson, H. H. (2018). Motor imagery involves predicting the sensory consequences of the imagined movement. Nature Communications, 9(1), 1617. doi:10.1038/s41467-018-03989-0

Knoblich, G., & Kircher, T. T. (2004). Deceiving oneself about being in control: conscious detection of changes in visuomotor coupling. Journal of Experimental Psychology: Human Perception and Performance, 30(4), 657–666. doi:10.1037/0096-1523.30.4.657

Kording, K. P., & Wolpert, D. M. (2006). Bayesian decision theory in sensorimotor control. Trends in Cognitive Sciences, 10(7), 319–326. doi:10.1016/j.tics.2006.05.003

Kuhn, S., Brass, M., & Haggard, P. (2013). Feeling in control: Neural correlates of experience of agency. Cortex, 49(7), 1935–1942. doi:10.1016/j.cortex.2012.09.002

Kwisthout, J., Bekkering, H., & van Rooij, I. (2017). To be precise, the details don’t matter: On predictive processing, precision, and level of detail of predictions. Brain and Cognition, 112, 84–91. doi:10.1016/j.bandc.2016.02.008

Lalanne, C., & Lorenceau, J. (2004). Crossmodal integration for perception and action. Journal of Physiology, Paris, 98(1-3), 265–279. doi:10.1016/j.jphysparis.2004.06.001

Lawson, R. P., Rees, G., & Friston, K. J. (2014). An aberrant precision account of autism. Frontiers in Human Neuroscience, 8, 302. doi:10.3389/fnhum.2014.00302

Legrand, D. (2007). Pre-reflective self-as-subject from experiential and empirical perspectives. Consciousness and Cognition, 16(3), 583–599. doi:10.1016/j.concog.2007.04.002

Lochmann, T., & Deneve, S. (2011). Neural processing as causal inference. Current Opinion in Neurobiology, 21(5), 774–781. doi:10.1016/j.conb.2011.05.018

Maglio, S. J., & Kwok, C. Y. (2016). Anticipated ambiguity prolongs the present: Evidence of a return trip effect. Journal of Experimental Psychology: General, 145(11), 1415–1419. doi:10.1037/xge0000228

Makwana, M., & Srinivasan, N. (2017). Intended outcome expands in time. Scientific Reports, 7(1), 6305. doi:10.1038/s41598-017-05803-1

McIntyre, J., Zago, M., Berthoz, A., & Lacquaniti, F. (2001). Does the brain model Newton’s laws? Nature Neuroscience, 4(7), 693–694. doi:10.1038/89477

McNally, R. J., Mair, P., Mugno, B. L., & Riemann, B. C. (2017). Co-morbid obsessive-compulsive disorder and depression: a Bayesian network approach. Psychological Medicine, 47(7), 1204–1214. doi:10.1017/S0033291716003287

Meder, B., Mayrhofer, R., & Waldmann, M. R. (2014). Structure induction in diagnostic causal reasoning. Psychological Review, 121(3), 277–301. doi:10.1037/a0035944

Miall, R. C., & Wolpert, D. M. (1996). Forward models for physiological motor control. Neural Networks, 9(8), 1265–1279. doi:10.1016/s0893-6080(96)00035-4

Mifsud, N. G., & Whitford, T. J. (2017). Sensory attenuation of self-initiated sounds maps onto habitual associations between motor action and sound. Neuropsychologia, 103, 38–43. doi:10.1016/j.neuropsychologia.2017.07.019

Miyazaki, M., & Hiraki, K. (2006). Delayed intermodal contingency affects young children’s recognition of their current self. Child Development, 77(3), 736–750. doi:10.1111/j.1467-8624.2006.00900.x

Montero, P., & Vilar, J. A. (2014). TSclust: An R package for time series clustering. Journal of Statistical Software, 62(1), 1–43. doi:10.18637/jss.v062.i01

Moore, J. W., & Fletcher, P. C. (2012). Sense of agency in health and disease: a review of cue integration approaches. Consciousness and Cognition, 21(1), 59–68. doi:10.1016/j.concog.2011.08.010

Moore, J. W., & Haggard, P. (2008). Awareness of action: Inference and prediction. Consciousness and Cognition, 17(1), 136–144. doi:10.1016/j.concog.2006.12.004

Moore, J. W., Lagnado, D., Deal, D. C., & Haggard, P. (2009). Feelings of control: contingency determines experience of action. Cognition, 110(2), 279–283. doi:10.1016/j.cognition.2008.11.006

Moore, J. W., & Obhi, S. S. (2012). Intentional binding and the sense of agency: a review. Consciousness and Cognition, 21(1), 546–561. doi:10.1016/j.concog.2011.12.002

Moreno-Bote, R., Knill, D. C., & Pouget, A. (2011). Bayesian sampling in visual perception. Proceedings of the National Academy of Sciences of the United States of America, 108(30), 12491–12496. doi:10.1073/pnas.1101430108

Moutoussis, M., Fearon, P., El-Deredy, W., Dolan, R. J., & Friston, K. J. (2014). Bayesian inferences about the self (and others): a review. Consciousness and Cognition, 25, 67–76. doi:10.1016/j.concog.2014.01.009

Park, J., Schlag-Rey, M., & Schlag, J. (2003). Voluntary action expands perceived duration of its sensory consequence. Experimental Brain Research, 149(4), 527–529. doi:10.1007/s00221-003-1376-x

Picard, F., & Friston, K. (2014). Predictions, perception, and a sense of self. Neurology, 83(12), 1112–1118. doi:10.1212/WNL.0000000000000798

Poulet, J. F., & Hedwig, B. (2002). A corollary discharge maintains auditory sensitivity during sound production. Nature, 418(6900), 872–876. doi:10.1038/nature00919

Powers, A. R., Mathys, C., & Corlett, P. R. (2017). Pavlovian conditioning-induced hallucinations result from overweighting of perceptual priors. Science, 357(6351), 596–600. doi:10.1126/science.aan3458

Press, C., Berlot, E., Bird, G., Ivry, R., & Cook, R. (2014). Moving time: the influence of action on duration perception. Journal of Experimental Psychology: General, 143(5), 1787–1793. doi:10.1037/a0037650

R Core Team. (2017). R: A language and environment for statistical computing. Retrieved from https://www.r- project.org/

Rao, R. P., & Ballard, D. H. (1999). Predictive coding in the visual cortex: a functional interpretation of some extra-classical receptive-field effects. Nature Neuroscience, 2(1), 79–87. doi:10.1038/4580

Repp, B. H., & Su, Y. H. (2013). Sensorimotor synchronization: a review of recent research (2006-2012). Psychonomic Bulletin & Review, 20(3), 403–452. doi:10.3758/s13423-012-0371-2

Ritterband-Rosenbaum, A., Nielsen, J. B., & Christensen, M. S. (2014). Sense of agency is related to gamma band coupling in an inferior parietal-preSMA circuitry. Frontiers in Human Neuroscience,8, 510. doi:10.3389/fnhum.2014.00510

Ruess, M., Thomaschke, R., & Kiesel, A. (2017). The time course of intentional binding. Attention, Perception, & Psychophysics, 79(4), 1123–1131. doi:10.3758/s13414-017-1292-y

Ruess, M., Thomaschke, R., & Kiesel, A. (2018). Intentional binding of visual effects. Attention, Perception, & Psychophysics, 80(3), 713–722. doi:10.3758/s13414-017-1479-2

Sato, A., & Yasuda, A. (2005). Illusion of sense of self-agency: discrepancy between the predicted and actual sensory consequences of actions modulates the sense of self-agency, but not the sense of self-ownership. Cognition, 94(3), 241–255. doi:10.1016/j.cognition.2004.04.003

Schwarz, K. A., Pfister, R., Kluge, M., Weller, L., & Kunde, W. (2018). Do we see it or not? Sensory attenuation in the visual domain. Journal of Experimental Psychology: General, 147(3), 418–430. doi:10.1037/xge0000353

Scutari, M. (2010). Learning Bayesian networks with the bnlearn R package. Journal of Statistical Software, 35(3), 1–22. doi:10.18637/jss.v035.i03

Sharps, M. J., & Martin, S. S. (2002). “Mindless” decision making as a failure of contextual reasoning. Journal of Psychology, 136(3), 272–282. doi:10.1080/00223980209604155

Shipp, S., Adams, R. A., & Friston, K. J. (2013). Reflections on agranular architecture: predictive coding in the motor cortex. Trends in Neurosciences, 36(12), 706–716. doi:10.1016/j.tins.2013.09.004

Stahl, A. E., & Feigenson, L. (2015). Cognitive development. Observing the unexpected enhances infants’ learning and exploration. Science, 348(6230), 91–94. doi:10.1126/science.aaa3799

Statisticat LLC. (2016). LaplacesDemon: Complete environment for Bayesian inference. R package version 16.1.0. Retrieved from https://cran.r-project.org/package=LaplacesDemon

Stetson, C., Cui, X., Montague, P. R., & Eagleman, D. M. (2006). Motor-sensory recalibration leads to an illusory reversal of action and sensation. Neuron, 51(5), 651–659. doi:10.1016/j.neuron.2006.08.006

Synofzik, M., Vosgerau, G., & Newen, A. (2008). Beyond the comparator model: a multifactorial two-step account of agency. Consciousness and Cognition, 17(1), 219–239. doi:10.1016/j.concog.2007.03.010

Takahata, K., Takahashi, H., Maeda, T., Umeda, S., Suhara, T., Mimura, M., & Kato, M. (2012). It’s not my fault: postdictive modulation of intentional binding by monetary gains and losses. PLoS ONE, 7(12), e53421. doi:10.1371/journal.pone.0053421

Teufel, C., Subramaniam, N., Dobler, V., Perez, J., Finnemann, J., Mehta, P. R., … Fletcher, P. C. (2015). Shift toward prior knowledge confers a perceptual advantage in early psychosis and psychosis-prone healthy individuals. Proceedings of the National Academy of Sciences of the United States of America, 112(43), 13401–13406. doi:10.1073/pnas.1503916112

Tomassini, A., Gori, M., Baud-Bovy, G., Sandini, G., & Morrone, M. C. (2014). Motor commands induce time compression for tactile stimuli. Journal of Neuroscience, 34(27), 9164–9172. doi:10.1523/JNEUROSCI.2782-13.2014

Treisman, M. (2013). The information-processing model of timing (Treisman, 1963): Its sources and further development. Timing & Time Perception, 1(2), 131–158. doi:10.1163/22134468-00002017

Uithol, S., & Paulus, M. (2014). What do infants understand of others’ action? A theoretical account of early social cognition. Psychological Research, 78(5), 609–622. doi:10.1007/s00426-013-0519-3

van de Ven, N., van Rijswijk, L., & Roy, M. M. (2011). The return trip effect: why the return trip often seems to take less time. Psychonomic Bulletin & Review, 18(5), 827–832. doi:10.3758/s13423-011-0150-5

Voss, M., Chambon, V., Wenke, D., Kuhn, S., & Haggard, P. (2017). In and out of control: brain mechanisms linking fluency of action selection to self-agency in patients with schizophrenia. Brain, 140(8), 2226–2239. doi:10.1093/brain/awx136

Walsh, E., & Haggard, P. (2013). Action, prediction, and temporal awareness. Acta Psychologica, 142(2), 220–229. doi:10.1016/j.actpsy.2012.11.014

Waszak, F., Cardoso-Leite, P., & Hughes, G. (2012). Action effect anticipation: neurophysiological basis and functional consequences. Neuroscience and Biobehavioral Reviews, 36(2), 943–959. doi:10.1016/j.neubiorev.2011.11.004

Weiss, C., Herwig, A., & Schutz-Bosbach, S. (2011). The self in action effects: selective attenuation of self-generated sounds. Cognition, 121(2), 207–218. doi:10.1016/j.cognition.2011.06.011

Wen, W., Brann, E., Di Costa, S., & Haggard, P. (2018). Enhanced perceptual processing of self-generated motion: Evidence from steady-state visual evoked potentials. NeuroImage, 175, 438–448. doi:10.1016/j.neuroimage.2018.04.019

Wenke, D., & Haggard, P. (2009). How voluntary actions modulate time perception. Experimental Brain Research, 196(3), 311–318. doi:10.1007/s00221-009-1848-8

Wickham, H. (2009). ggplot2: Elegant Graphics for Data Analysis. New York: Springer-Verlag

Wolpe, N., Haggard, P., Siebner, H. R., & Rowe, J. B. (2013). Cue integration and the perception of action in intentional binding. Experimental Brain Research, 229(3), 467–474. doi:10.1007/s00221-013-3419-2

Wolpe, N., Ingram, J. N., Tsvetanov, K. A., Geerligs, L., Kievit, R. A., Henson, R. N., … Rowe, J. B. (2016). Ageing increases reliance on sensorimotor prediction through structural and functional differences in frontostriatal circuits. Nature Communications, 7, 13034. doi:10.1038/ncomms13034

Wolpert, D. M. (1997). Computational approaches to motor control. Trends in Cognitive Sciences, 1(6), 209–216. doi:10.1016/S1364-6613(97)01070-X

Wolpert, D. M., Miall, R. C., & Kawato, M. (1998). Internal models in the cerebellum. Trends in Cognitive Sciences, 2(9), 338–347. doi:10.1016/S1364-6613(98)01221-2

Yarkoni, T., & Westfall, J. (2017). Choosing prediction over explanation in psychology: Lessons from machine learning. Perspectives on Psychological Science, 12(6), 1100–1122. doi:10.1177/1745691617693393

Yarrow, K., Haggard, P., Heal, R., Brown, P., & Rothwell, J. C. (2001). Illusory perceptions of space and time preserve cross-saccadic perceptual continuity. Nature, 414(6861), 302–305. doi:10.1038/35104551

Yoshie, M., & Haggard, P. (2013). Negative emotional outcomes attenuate sense of agency over voluntary actions. Current Biology, 23(20), 2028–2032. doi:10.1016/j.cub.2013.08.034

Yoshie, M., & Haggard, P. (2017). Effects of emotional valence on sense of agency require a predictive model. Scientific Reports, 7(1), 8733. doi:10.1038/s41598-017-08803-3

Zhao, K., Chen, Y. H., Yan, W. J., & Fu, X. (2013). To bind or not to bind? Different temporal binding effects from voluntary pressing and releasing actions. PLoS ONE, 8(5), e64819. doi:10.1371/journal.pone.0064819

